# Four enzymes control natural variation in the steroid core of *Erysimum* cardenolides

**DOI:** 10.1101/2024.04.10.588904

**Authors:** Gordon C. Younkin, Martin L. Alani, Tobias Züst, Georg Jander

## Abstract

Plants commonly produce families of structurally related metabolites with similar defensive functions. This apparent redundancy raises the question of underlying molecular mechanisms and adaptive benefits of such chemical variation. Cardenolides, a class defensive compounds found in the wallflower genus *Erysimum* (L., Brassicaceae) and scattered across other plant families, show substantial structural variation, with glycosylation and hydroxylation being common modifications of a steroid core, which itself may vary in terms of stereochemistry and saturation. Through a combination of chemical mutagenesis and analysis of gene coexpression networks, we identified four enzymes involved in cardenolide biosynthesis in *Erysimum* that work together to determine stereochemistry at carbon 5 of the steroid core: Ec3βHSD, a 3β-hydroxysteroid dehydrogenase, Ec3KSI, a ketosteroid isomerase, EcP5βR2, a progesterone 5β-reductase, and EcDET2, a steroid 5α-reductase. We biochemically characterized the activity of these enzymes *in vitro* and generated CRISPR/Cas9 knockout lines to confirm activity *in vivo*. Cardenolide biosynthesis was not eliminated in any of the knockouts. Instead, mutant plants accumulated cardenolides with altered saturation and stereochemistry of the steroid core. Furthermore, we found variation in carbon 5 configuration among the cardenolides of 44 species of *Erysimum*, where the occurrence of some 5β-cardenolides is associated with the expression and sequence of P5βR2. This may have allowed *Erysimum* species to fine-tune their defensive profiles to target specific herbivore populations over the course of evolution.

**SIGNIFICANCE STATEMENT:** Plants use an array of toxic compounds to defend themselves from attack against insects and other herbivores. One mechanism through which plants may evolve more toxic compounds is through modifications to the structure of compounds they already produce. In this study, we show how plants in the wallflower genus *Erysimum* use four enzymes to fine-tune the structure of toxic metabolites called cardenolides. Natural variation in the sequence and expression of a single enzyme called progesterone 5β-reductase 2 partly explains the variation in cardenolides observed across the *Erysimum* genus. These alterations to cardenolide structure over the course of evolution suggests that there may be context-dependent benefits to *Erysimum* to invest in one cardenolide variant over another.

## INTRODUCTION

It is well established that plants synthetize specialized compounds with toxic properties to defend themselves from herbivore attack (1–3), with the presence or absence of specific compounds shaping herbivore community structure (4). However, due to physiological and phylogenetic constraints, plants usually only produce a few, structurally related toxic compounds. Because of this limitation, structural diversity within metabolite classes and modifications to existing metabolites play an essential role in regulating herbivore resistance. In fact, there is substantial evidence that small, biochemically accessible modifications to the structure of existing metabolites are a critical mechanism through which plants evolve new, more potent defenses in response to herbivore pressure. For example, in the furanocoumarin-producing lineages of the Apiaceae, a single enzyme transforms umbelliferone to xanthotoxin, which is far more toxic and restricted in its occurrence (5). An analysis of *Inga* foliar metabolomes proposed a model where differences in chemical diversity between species were best explained by regulatory changes to enzymes in existing biosynthetic pathways, allowing plants to rapidly alter their chemical defenses during evolution, facilitating adaptation and divergence between closely related plant lineages (6).

Cardenolides, which inhibit Na^+^/K^+^-ATPases in animals and evolved repeatedly in diverse plant lineages (7), exhibit variation in the stereochemical configuration of carbon 5 of their steroid core (Figure 1). Fixation of carbon 5 configuration likely occurs during a series of steps involving repeated oxidation and reduction of pregnane intermediates. In plants, as in animals, such reactions are mostly catalyzed by hydroxysteroid dehydrogenases, either from the short chain dehydrogenase/reductase (SDR) or the aldo-keto reductase (AKR) families (8–10), though progesterone-5β-reductases (P5βRs) also play a role in plants (11–13).

**Figure 1.**
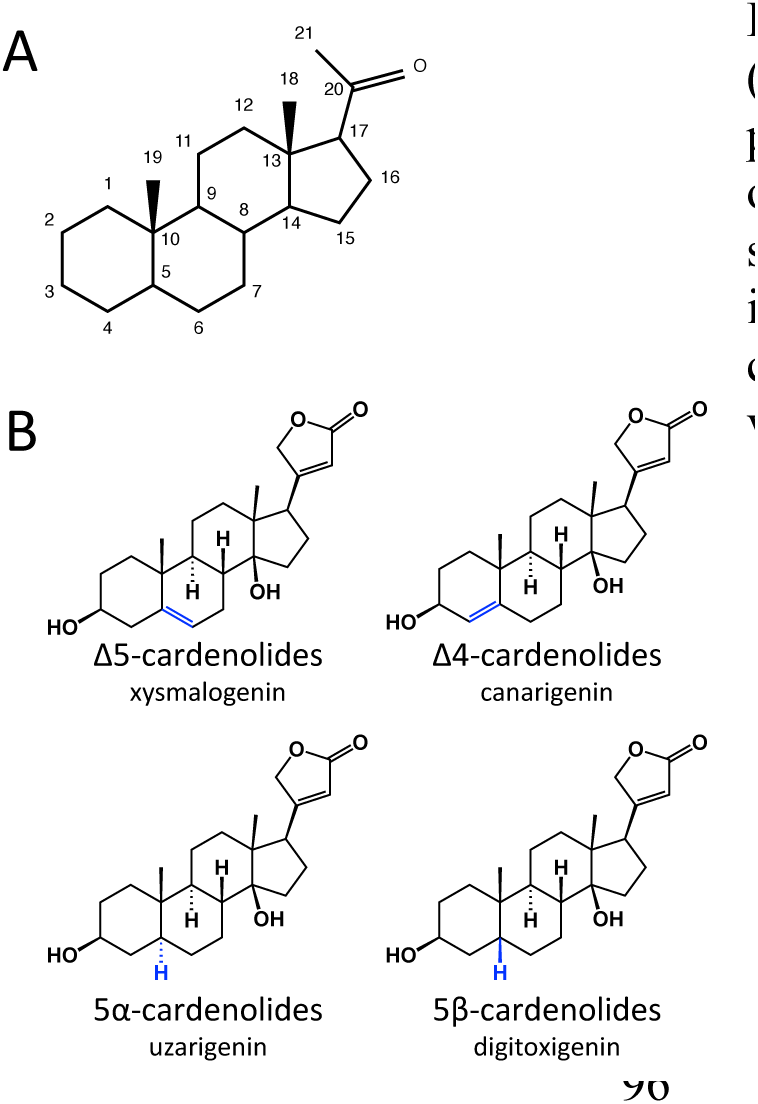
Variation in the cardenolide steroid core. (A) Carbon numbering scheme for the steroid core of pregnanes and cardenolides. (B) Four structural classes of cardenolides. Variable position of double bond or stereochemical configuration of carbon 5 is highlighted in blue. One example of each of the four structural classes of cardenolides is shown. They may be modified via hydroxylation, glycosylation, etc.

The relative activity of these enzymes on cardenolide intermediates may help explain the natural variation seen in cardenolide configuration at carbon 5 across plant lineages. For example, 5β-cardenolides are commonly found in *Digitalis* (Plantaginaceae) (14), whereas many members of the Apocynaceae family accumulate 5α-cardenolides (15). Less commonly reported are unsaturated cardenolides such as Δ5,6-unsaturated xysmalogenin and Δ4,5-unsaturated canarigenin (Figure 1B). Xysmalogenin has been identified in some members of the Apocynaceae, including *Gomphocarpus sinaicus* Boiss. (16), *Asclepias curassavica* L. (17), *Periploca sepium* Bunge (18), as well as in *Isoplexis* spp. (Plantaginaceae, sometimes classified *Digitalis*). (19). Canarigenin has also been reported from *Isoplexis* spp. (19, 20) and *Convallaria majalis* L. (Asparagaceae) (21). In *Erysimum*, both 5α- and 5β-cardenolides occur across the genus, including co-occurrence of both in many species (22, 23). At least one species, *Erysimum x allionii* (formerly *Cheiranthus allionii*), also accumulates Δ4-cardenolides (24). The functional implications of this structural variation at carbon 5 has been the subject of some research in both medical and ecological contexts, with substantial impact on target-site binding, inhibitory activity, and toxicity. However, much remains unclear in this regard (25–27).

Enzymes such as hydroxylases and glycosyltransferases that modify pathway end products are often cited as a driver of structural diversity within classes of defensive metabolites, *e.g.*, in the well-studied glucosinolate and benzoxazinoid pathways (28, 29). By contrast, we provide an example of core pathway enzymes that control structural diversity of pathway end products. Specifically, we show that the structure of cardenolides produced by the genus *Erysimum* depends on the sequence and expression of four key enzymes influencing the stereochemistry and saturation at carbon 5 of the steroid core.

## RESULTS

### Mutant screens

Two independent ethyl methanesulfonate (EMS)-mutagenized lines with altered cardenolide content were characterized, and the causal mutations identified via bulked segregant analysis (BSA). EMS mutant line #635 accumulates low levels of the cardenolides found in wildtype *E. cheiranthoides* (Figure 2A; Table S1) and instead accumulates peaks with cardenolide-like fragmentation and a *m/z* two Daltons less than digitoxigenin glycosides, suggesting that they may be Δ4-or Δ5-unsaturated cardenolides (Figures 1B). This phenotype mapped to a region on chromosome 7 (Figure 2B) containing a G148E missense mutation in *Erche07g001535* (*Ec3βHSD*), a gene encoding a 3β-hydroxy-Δ5-steroid dehydrogenase (3βHSD) (Figure 2C).

**Figure 2.**
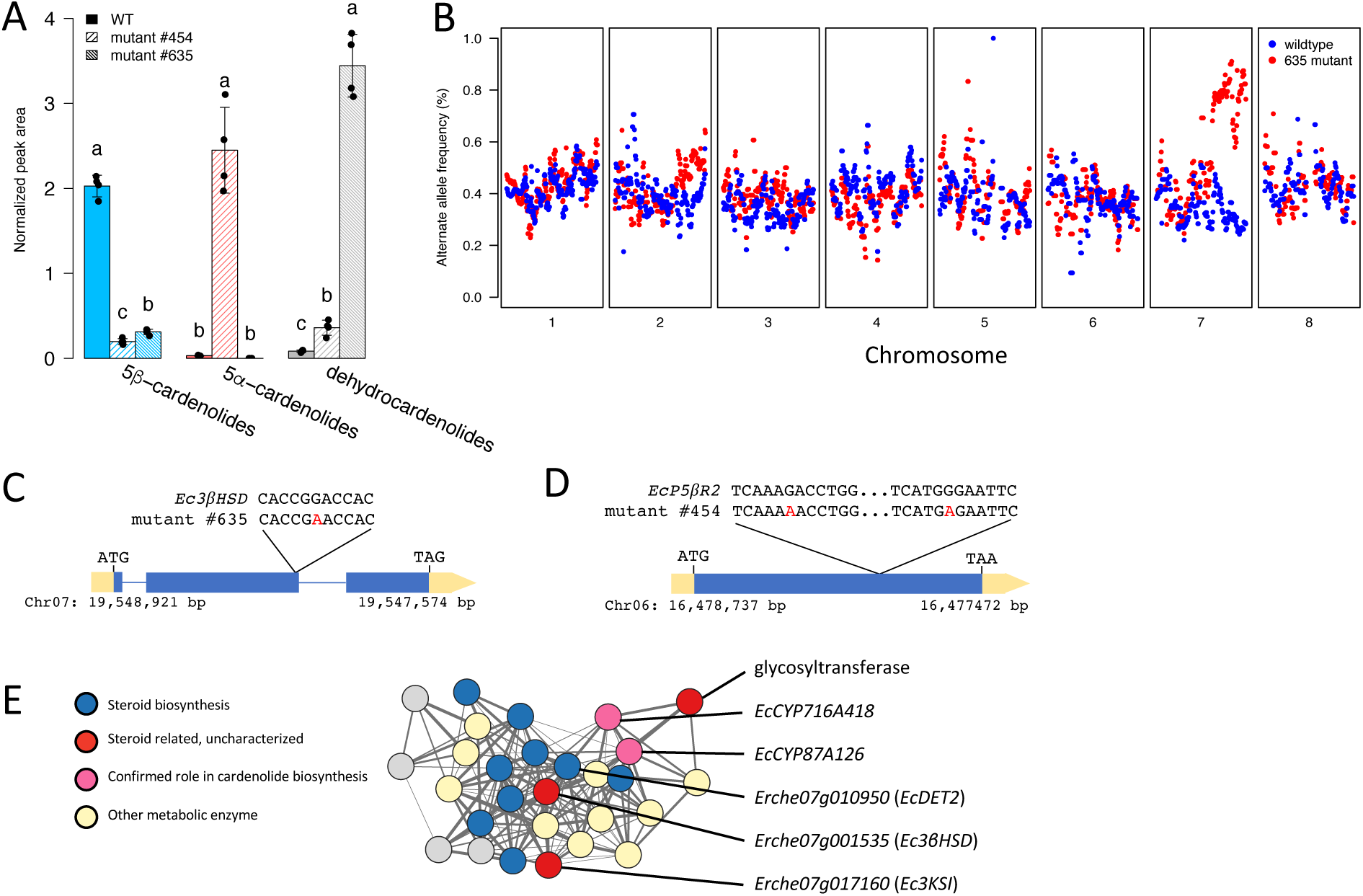
Identification of candidate genes for cardenolide biosynthesis in *Erysimum cheiranthoides*. (A) Cardenolide abundances in wildtype (WT) *E. cheiranthoides* and ethyl methanesulfonate (EMS) mutant lines. Cardenolides in mutant #454 have the same mass as cardenolides found in WT plants but elute at different retention times. Presumed dehydrocardenolides are identified by a characteristic genin at *m/z*=373.2379. N = 4, error bars indicate ± s.d., letters are p<0.001, ANOVA with post-hoc Tukey’s HSD on log-transformed peak areas. (B) Mapping results of bulked segregant analysis for mutant #635. Alternate (mutant) allele frequency, smoothed over 1 Mbp segments, is plotted across the eight *E. cheiranthoides* chromosomes. In plants with a mutant chemotype (red), mutant alleles dominate in the latter half of chromosome seven. (C) Within this region is a 3β-hydroxysteroid dehydrogenase (*Ec3βHSD*) with a G148E missense mutation in mutant #635 plants. (D) In mutant #454 plants, there are two EMS-induced mutations in the coding region of *EcP5βR2.* For mapping results for mutant #454, see Figure S1. Exons are shown as blue rectangles, introns as blue lines, and untranslated regions as yellow rectangles. (E) Cardenolide-related gene coexpression cluster from an analysis of transcript abundances in 48 *Erysimum* species. *EcCYP716A418* and *EcCYP87A126* are cytochrome P450s involved in cardenolide biosynthesis in *E. cheiranthoides*. Other candidates for involvement in cardenolide biosynthesis include a 5α-reductase (*EcDET2*), *Ec3βHSD*, and a 3-ketosteroid isomerase (*Ec3KSI*).

Enzymes of this type are thought to be required for cardenolide biosynthesis (30–32).

EMS mutant #454 was described by Mirzaei et al. (33), with the cardenolide phenotype being linked to a locus on chromosome 6 (Figure S1). However, no causal mutation was identified in that study. We re-evaluated the cardenolide phenotype and found that mutant #454 plants accumulate compounds with the same mass as digitoxigenin glycosides, but with shifted retention times (Figure S2). We hypothesized that these peaks represent uzarigenin glycosides with 5α stereochemistry (Figure 1B, 2A; Table S1). This led us to the identification of *Erche06g007150* (*EcP5βR2*), which encodes a progesterone-5β-reductase (P5βR). This group of enzymes catalyzes the stereospecific reduction of α,β-unsaturated ketones including progesterone, methylvinylketone, and 2-cyclohexene-1-one (34). *EcP5βR2* contained two amino acid mutations (R184K and G201R) at the genetically linked locus on chromosome 6 (Figure 2D). We therefore hypothesized that in the absence of a functional P5βR, a steroid 5α-reductase (5αR) acts on progesterone, resulting in the accumulation of 5α-cardenolides.

### Gene coexpression analysis

Gene coexpression analysis across 48 *Erysimum* species revealed a cluster of 28 coexpressed genes related to steroid metabolism and cardenolide biosynthesis (Figure 2E). Two genes, *EcCYP87A126* and *EcCYP716A418*, are involved in cardenolide biosynthesis (35), and one, *Ec3βHSD*, was identified in EMS mutant #635. Eight genes in the cluster encode enzymes that are directly involved in core sterol or isoprenoid metabolism (Table S2). The remaining genes are considered candidates for involvement in cardenolide biosynthesis. Of note is a gene encoding a short-chain dehydrogenase/reductase (SDR), *Erche07g017160* (*Ec3KSI*). The closest Arabidopsis ortholog, *AT2G33630*, is annotated as having a 3β-hydroxysteroid-dehydrogenase/isomerase domain (IPR002225)(36). An additional candidate from the coexpression cluster is a steroid 5αR, *Erche07g010950* (*EcDET2*), which is involved in brassinosteroid biosynthesis (37) and may also be involved in 5α-cardenolide biosynthesis.

### Functional characterization of candidate enzymes

We examined the role of candidate enzymes for involvement in cardenolide biosynthesis by functionally characterizing the recombinant purified proteins *in vitro*. Recombinant Ec3βHSD had steroid-3-dehydrogenase activity on pregnenolone **2** to form isoprogesterone **3** in the presence of NAD+. We also saw some production of progesterone **4**, implying that the enzyme may additionally have Δ5,4 isomerase activity (Figure 3A, S3; Table S3). As plant 3βHSD enzymes typically do not possess Δ5,4 isomerase activity (31, 38, 39), we independently checked for isomerase activity by supplying the enzyme directly with isoprogesterone **3**, but we did not see an increase in isomerization to progesterone **4** relative to a negative control (Figure 3B, S3; Table S3).

**Figure 3.**
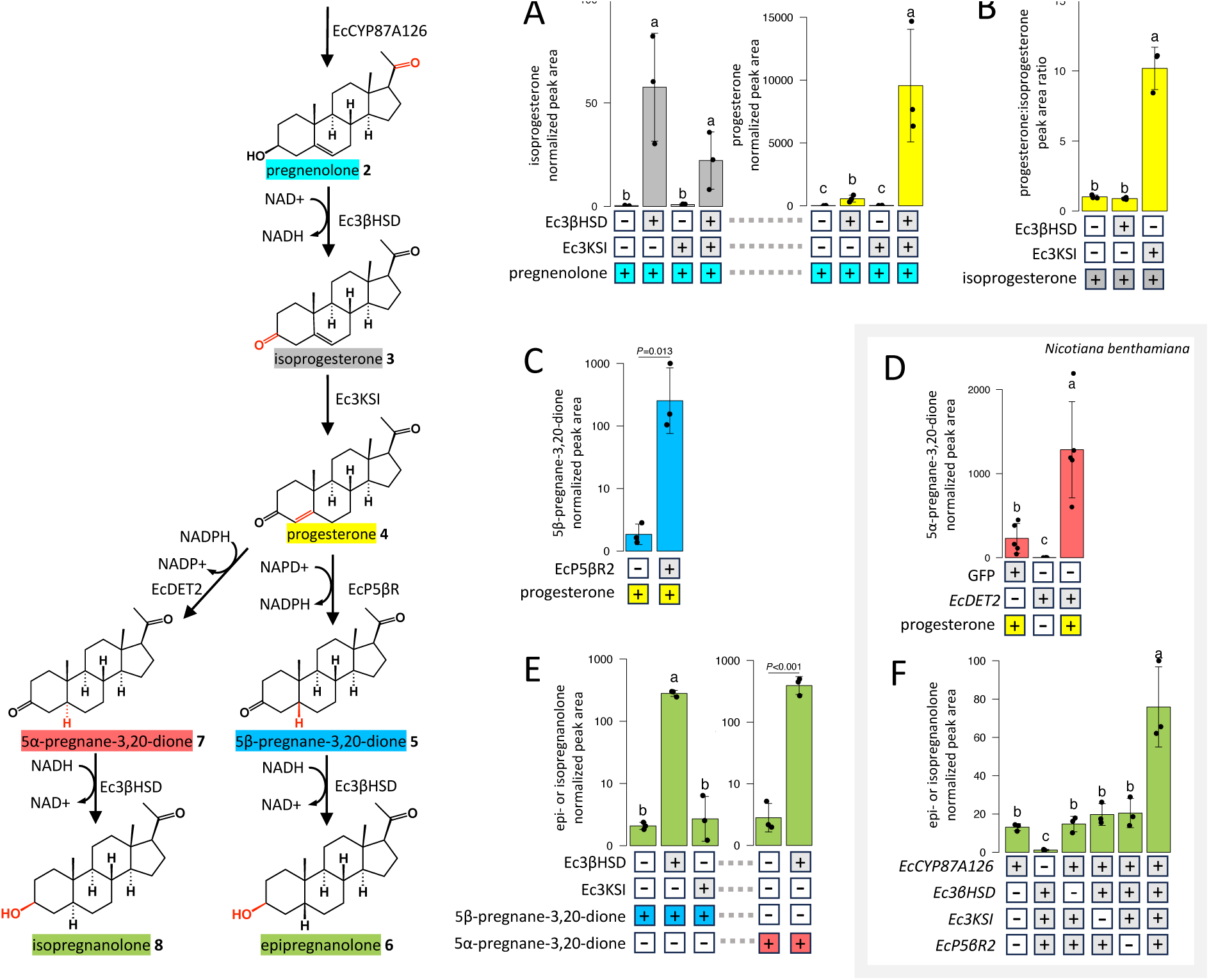
Characterization of candidate cardenolide biosynthetic enzymes from *Erysimum cheiranthoides.* Assays are *in vitro* with purified recombinant enzymes except where noted. (A) Conversion of pregnenolone to isoprogesterone and progesterone by 3β-hydroxysteroid dehydrogenase (Ec3βHSD) and 3-ketosteroid isomerase (Ec3KSI). (B) Conversion of isoprogesterone to progesterone by Ec3βHSD and Ec3KSI. Under assay conditions, ∼50% of the peak area is attributable to progesterone even in the negative control. (C) Conversion of progesterone to 5β-pregnane-3,20-dione by progesterone 5β-reductase 2 (*Ec*P5βR2). (D) In *N. benthamiana*, significantly more 5α-pregnane-3,20-dione is produced when progesterone is coinfiltrated with 5α-reductase (*EcDET2*) compared to a negative control. (E) Reduction of 5β-pregnane-3,20-dione and 5ɑ-pregnane-3,20-dione to epipregnanolone or a stereoisomer by Ec3βHSD but not Ec3KSI. (F) Production of epipregnanolone or stereoisomer when *EcCYP87A126*, *Ec3βHSD*, *Ec3KSI*, and *EcP5BR2* are coexpressed in *N. benthamiana.* For all assays: N = 3 replicates per enzyme, except in (D) where N=5. Error bars indicate ± s.d., letters are *P*<0.001, ANOVA with post-hoc Tukey’s HSD. *P*-values above bars are from Student’s *t*-test. Statistics were performed on log-transformed peak areas. Negative controls used a purified recombinant 2-oxoglutarate dioxygenase not otherwise discussed in this study. LCMS chromatograms and MSMS spectra associated with all assays are provided in Figure S3.

We also tested Ec3βHSD for the ability to catalyze the reverse reaction, the reduction of the 3-keto group of 5β-pregnane-3,20-dione **5** or 5α-pregnane-3,20-dione **7** to the 3β-hydroxyl in epipregnanolone **6** or isopregnanolone **8**. When Ec3βHSD was supplied with 5β-pregnane-3,20-dione **5** or 5α-pregnane-3,20-dione **7** as a substrate and NADH as a cofactor, we saw formation of a product with *m/z*=319.2637 in both cases, which is consistent with epipregnanolone **6** and isopregnanolone **8**, but we were unable to separate these two products and the epipregnanolone standard chromatographically (Figure 3E, S3; Table S3). Therefore, the exact stereochemical configuration of these predicted products is not confirmed.

Recombinant Ec3KSI converts isoprogesterone **3** to progesterone **4** *in vitro* (Figure 3B, S3). We also tested Ec3KSI for 3βHSD activity, but no activity was observed for either the oxidation or reduction reaction (Figure 3A,E, S3; Table S3). Furthermore, when we combined Ec3βHSD and Ec3KSI in a single reaction and supplied pregnenolone and NAD+, we observed consumption of isoprogesterone and increased formation of progesterone relative to the same reaction containing only Ec3βHSD (Figure 3A; Table S3). We therefore identified Ec3KSI as a 3-ketosteroid isomerase and showed that it works in concert with Ec3βHSD to convert pregnenolone **2** to progesterone **4**. When supplied with progesterone **4** and NADPH, recombinant EcP5βR2 catalyzes the formation of 5β-pregnane-3,20-dione **5** (Figure 3C, S3; Table S3).

EcDET2 is membrane-bound, complicating purification of the recombinant protein. We therefore coinfiltrated *EcDET2* with progesterone in *Nicotiana benthamiana* leaves. Although the 5α-reduction of progesterone **4** is catalyzed by endogenous enzymes in *N. benthamiana* leaves even in the GFP control, more 5α-pregnane-3,20-dione **7** is produced when EcDET2 is present, consistent with previous studies showing that DET2 orthologs can use progesterone **4** as a substrate (40) (Figure 3D, S3; Table S3).

### Production of epipregnanolone in N. benthamiana

To test whether the identified enzymes can work together *in planta* to produce intermediates in cardenolide biosynthesis, we coexpressed the sterol side chain cleaving enzyme that initiates cardenolide biosynthesis, EcCYP87A126 (35, 41) together with Ec3βHSD, Ec3KSI, and EcP5βR2 in leaves of *N. benthamiana* and observed the production of a compound with the same *m/z* and retention time as epipregnanolone **6**, although we cannot rule out that it may be a stereoisomer (Figure S3). A small amount of epipregnanolone **6** was detected as long as EcCYP87A126 was present, but significantly more epipregnanolone **6** was produced if Ec3βHSD, Ec3KSI, and EcP5βR2 were all included (Figure 3F; Table S3).

### Knockouts of candidate genes have altered cardenolide profiles

To assess the role of candidate enzymes in cardenolide biosynthesis *in vivo*, we generated two independent CRISPR/Cas9 knockout lines for all candidate genes (Figures S4-S8 for sequences of mutants). With the exception of *det2* knockout lines, which had the characteristic dwarf phenotype of brassinosteroid biosynthetic mutants (37) (Figure 4A), none of the mutant lines displayed obvious growth phenotypes. We analyzed cardenolide profiles via UPLC-MS, using both methanolic extracts of intact cardenolides and cardenolide extracts subjected to mild acidic conditions, which resulted in hydrolysis of the sugar moieties. This allowed us to directly compare the cardenolide genins produced by each mutant line. For Δ4-cardenolides (canarigenin glycosides), a water molecule is eliminated under acidic conditions to produce 3,5-anhydroperiplogenin (Figure 4C)(19, 20), which we used as a proxy for Δ4-cardenolide abundance. While we did not have an authentic standard for uzarigenin, xysmalogenin, canarigenin, 3,5-anhydroperiplogenin, cannogenol, or cannogenin, MSMS spectra together with pathway logic allowed us to identify these compounds with reasonable confidence. To further confirm our identification of uzarigenin, we performed acid hydrolysis on cardenolide extracts from uzarigenin-containing *Calotropis procera* leaves (Figure S9)(42).

**Figure 4.**
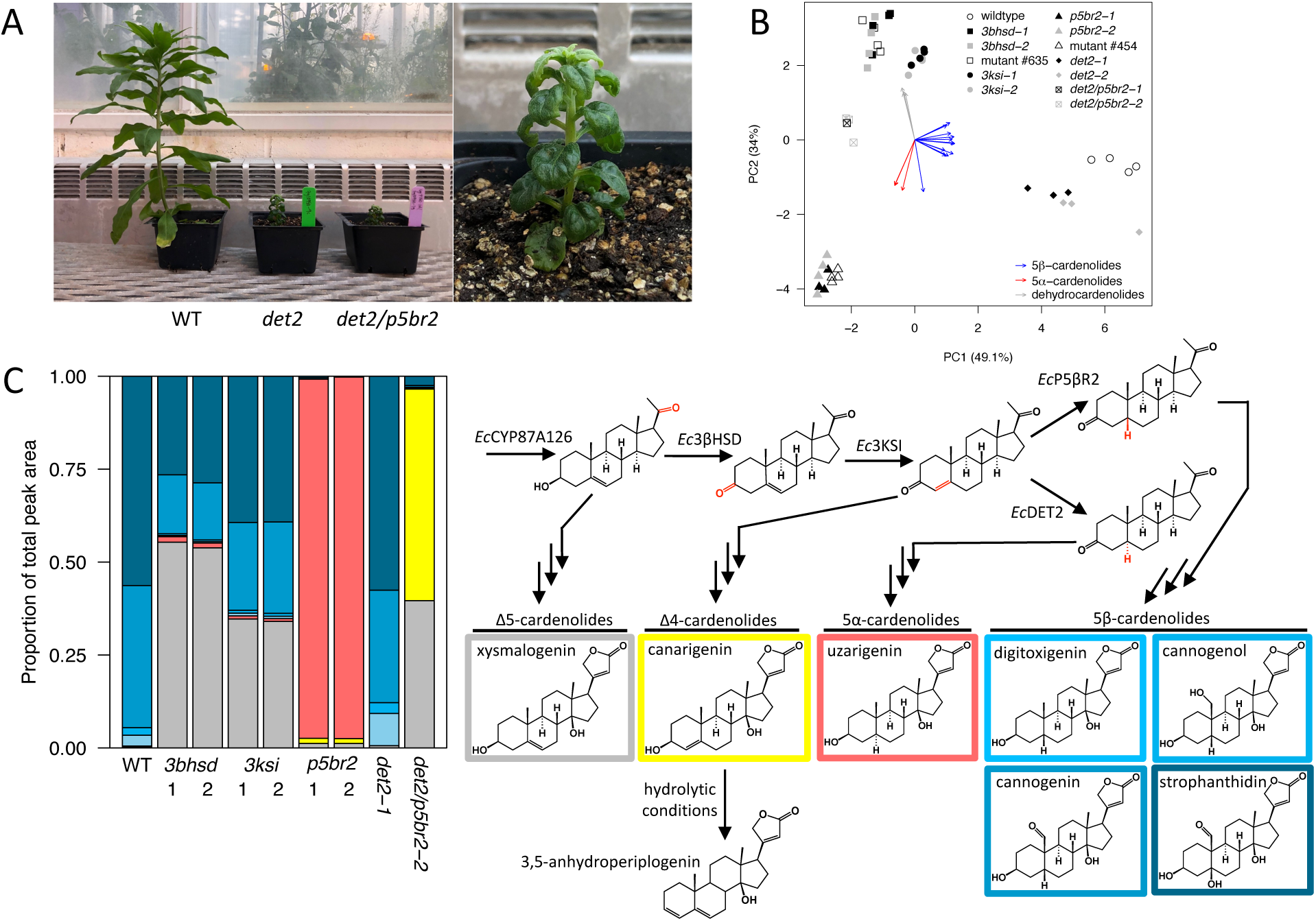
Characterization of *Erysimum cheiranthoides* cardenolide biosynthesis mutants. (A) Photo of *E. cheiranthoides* wildtype (WT), a steroid 5α-reductase (*det2*) knockout, and a DET2/progesterone 5β-reductase (*det2/p5br*) double mutant, which display a dwarf phenotype characteristic of brassinosteroid biosynthesis mutants. (B) Principal component analysis of intact cardenolides detected by UPLC-MS in *E. cheiranthoides* WT and mutant lines: 3β-hydroxysteroid dehydrogenase (*3bhsd*), 3-ketosteroid isomerase (*3ksi*). Ethyl methane-sulfonate (EMS) mutants #454 and #635 cluster with corresponding CRISPR/Cas9 mutant lines. Arrows indicate loadings for individual cardenolide mass features and are grouped into structural classes by color. (C) Relative peak area of cardenolide genins following acid hydrolysis of *E. cheiranthoides* mutant lines (N = 3 plants per line). For Δ5-cardenolides (grey; xysmalogenin), 5α-cardenolides (red; uzarigenin), and 5β-cardenolides (blue; digitoxigenin, cannogenin, cannogenol, and strophanthidin), the cardenolide genins were detected directly via UPLC-MS. Δ4-cardenolides undergo dehydration under acid conditions, so 3,5-anhydroperiplogenin was detected as a proxy for canarigenin (yellow).

Based on a principal component analysis (PCA) of intact cardenolides in the mutant lines, we confirmed a causal relationship between the genetically linked mutations in EMS mutants #454 and #635 and their cardenolide phenotypes (Figure 4B; Table S1). Hydrolyzed extracts of wildtype *E. cheiranthoides* and *det2* single mutants were dominated by the 5β-cardenolide series: digitoxigenin, cannogenol, cannogenin, and strophanthidin. All other mutant lines had altered cardenolide profiles relative to wildtype, but cardenolide production was not eliminated in any of the mutants. *3bhsd* and *3ksi* lines had similar cardenolide profiles, with both lines accumulating lower levels of 5β-cardenolides compared to wildtype and containing Δ5-cardenolides with *m/z*=373.2379, which are not found in wildtype *E. cheiranthoides* (Figure 4C, S10; Table S4). Although the chemotype of *3bhsd* and *ksi* lines was qualitatively very similar, the reduction in 5β-cardenolide abundance was more severe in *3bhsd* plants (Figure S11; Table S5). One possible explanation for this is a degree of functional overlap between these enzymes. To test whether Ec3βHSD and Ec3KSI have redundant roles in cardenolide biosynthesis, we crossed *3bhsd-1* and *3ksi-1* lines to generate *3bhsd*/*3ksi* double mutants. The double mutants had a cardenolide profile very similar to *3bhsd* plants (Figure S11; Table S5), suggesting that Ec3βHSD and Ec3KSI have distinct roles. If the two enzymes were redundant, we would expect an additive effect on the cardenolide phenotype in the double mutant.

Acid hydrolysis confirmed the digitoxigenin-glycoside isomers in *p5br2* plants to be uzarigenin glycosides, with 5α configuration (Figure 4C). Additionally, *p5br2* lines accumulated the same *m/z*=373.2379 peaks observed in *3bhsd* and *3ksi* lines, which we observed to be a mix of Δ4- and Δ5-cardenolides following hydrolysis (Figure 4C; Table S4). Notably, cardenolide hydroxylation (resulting in the derived cardenolide genins cannogenol, cannogenin, and strophanthidin) is eliminated in *p5br2* plants, suggesting that the cardenolide hydroxylases expressed in *E. cheiranthoides* act only on 5β-cardenolides (Figure 4C, S10). Although these results clearly demonstrate the involvement of EcP5βR2 in cardenolide biosynthesis, we also investigated a paralogous gene, *EcP5βR1* (*Erche02g027660*), for potential involvement in the pathway. EcP5βR1 acted on progesterone **4** to produce 5β-pregnane-3,20-dione **5** when co-infiltrated in *N. benthamiana* (Figure S12). However, in a EcP5βR1 knockout line, the cardenolide profile was unchanged (Figure S13; Table S6).

In order to test whether 5α-cardenolide production in *p5br2* mutants was mediated by EcDET2, we generated CRISPR/Cas9 knockouts of *Ec*DET2 in the *p5br2-1* background. In *p5br2*/*det2* double mutants, production of cardenolides with a fully reduced steroid core was nearly eliminated, and was replaced by accumulation of a mix of Δ4- and Δ5-cardenolides (Figure 4C, Table S4), confirming that EcDET2 acts as a 5αR in cardenolide biosynthesis in the absence of a functional EcP5βR2.

### Natural variation in progesterone 5β-reductase activity across the genus Erysimum

We conducted a survey of the cardenolide genins across the genus by subjecting methanolic leaf extracts from 44 species of *Erysimum* to acid hydrolysis. The following cardenolide genins were detected: the 5β-cardenolides digitoxigenin, cannogenol, cannogenin, and strophanthidin; the 5α-cardenolide uzarigenin; the Δ4-cardenolide canarigenin; the Δ5-cardenolide xysmalogenin; and an isomer of cannogenol that we speculate may be its 5α-isomer. With the exception of *E. collinum*, which does not produce detectable levels of cardenolides, both uzarigenin and strophanthidin were detected in hydrolyzed extracts of all species. Digitoxigenin, cannogenol, and cannogenin were relatively rare, being mostly restricted to the monophyletic clade containing *E. cheiranthoides*, *E. sylvestre*, and two closely related species of uncertain taxonomic identity (43)(Figure 5A). 3,5-anhydroperiplogenin (canarigenin) and xysmalogenin were also detected at low levels in some species (Figure 5A; Table S7).

**Figure 5.**
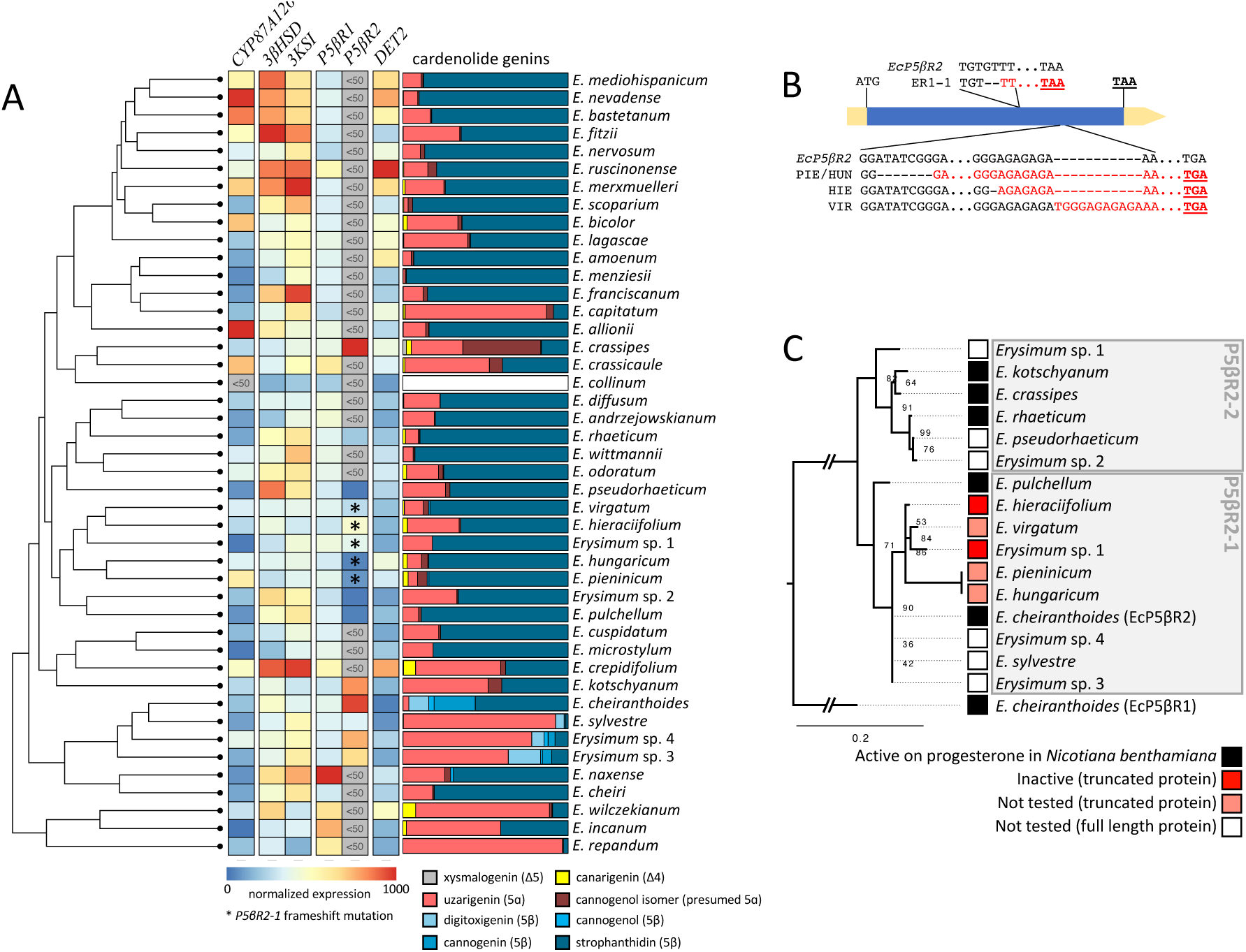
Gene expression and cardenolide configuration across 44 species of *Erysimum*. (A) Expression of cardenolide biosynthesis genes and detection of eight cardenolide genins following acid hydrolysis mapped against an *Erysimum* species phylogeny (Züst *et al* 2020).

Grey indicates very low expression (<50 counts per million reads). Expression for each gene is normalized separately. * indicates a frameshift mutation in *P5BR2-1*. Cardenolide genin abundances are displayed as percent of total LC-MS peak area of all genins, mean of n=1-3 replicates per species. No data are displayed for *E. collinum* due to very low levels of cardenolides. (B) Location and sequence of frame-shift mutations in the coding region of *EcP5ýR2* orthologs from *Erysimum* sp. 1 (ER1), *E. pieninicum* (PIE), *E. hungaricum* (HUN), *E. hieraciifolium* (HIE), and *E. virgatum* (VIR). Regions translated out of frame are indicated in red, with in-frame stop codons bold and underlined. (C) Amino acid phylogeny of *Erysimum* P5ýR2 proteins for which a full sequence could be recovered from the transcriptomes. Activity of selected enzymes were tested via transient expression and co-infiltration of progesterone in *Nicotiana benthamiana* leaves. Numbers at nodes indicate bootstrap support from 10,000 replicates and scale bar indicates estimated substitutions per site. Multiple sequence alignment underlying P5ýR2 protein phylogeny is available in Figure S14.

We next examined *P5βR2* and *DET2* sequences and expression levels across the *Erysimum* genus to better understand how they interact to determine relative levels of 5α- and 5β-cardenolides. Of the three *P5βR* sequences found in the *E. cheiranthoides* genome, only two have orthologs that are expressed in the species included in this study. *EcP5βR1* orthologs are uniformly expressed across the genus, but based on the *E. cheiranthoides p5br1* knockouts, they are unlikely to be involved in 5β-cardenolide biosynthesis. By contrast, *EcP5βR2* orthologs are only expressed at greater than 50 counts per million reads (CPM) in 15 of the 44 species examined (Figure 5A; Table S8), and in five of these species, *P5βR2* contains a frameshift mutation (Figure 5B, S14). Based on protein phylogeny, *Erysimum* P5βR2 proteins can be further classified into two clades, P5βR2-1 and P5βR2-2, with EcP5βR2 belonging to the P5βR2-1 clade (Figure 5C). We cloned *P5βR2* orthologs in both clades from six *Erysimum* species, as well as *EcP5βR1*, and assessed activity via co-infiltration with progesterone **4** in *N. benthamiana*. All full-length P5βR2 proteins were capable of converting progesterone **4** to 5β-pregnane-3,20-dione **5** *in planta*, while truncations resulted in a loss of activity (Figure 5C, S12). *Erysimum* sp. 1 was the only species to express both *P5βR2-1* and *P5βR2-2*, but the expressed *P5βR2-1* encodes a non-functional protein. Among species examined, the expression of a functional P5βR2-1 was required for production of the 5β-cardenolides digitoxigenin and cannogenin.

Orthologs of *Ec3βHSD*, *EcKSI*, and *EcDET2* are expressed across the genus, including in *Erysimum collinum*, where *CYP87A126*, the first gene in the pathway, is not expressed and very low levels of cardenolides are produced (43) (Figure 5A), suggesting that they may have roles in steroid metabolism outside of cardenolide biosynthesis. Close examination of the transcriptome data revealed that some species express more than one distinct transcript of the genes examined here, despite the *E. cheiranthoides* genome containing only one copy in the case of *Ec3βHSD* and *EcDET2*. However, the nature of transcriptomic data makes it difficult to assess genomic copy number, and it is unclear whether transcriptomic sequence variation is due to polypoidy in some species, gene duplication and sequence divergence, or allelic variation of a single locus.

### Phylogenetic analysis of E. cheiranthoides cardenolide biosynthesis genes

We inferred phylogenetic trees for *Ec3βHSD*, *EcKSI*, *EcP5βR*, and *EcDET2* to better understand their relationship to characterized genes from other species. All species examined had more than one *Ec3βHSD*-like gene, with the exception of *E. cheiranthoides*. The three orthologs from *E. crepidifolium* (44), two from *D. lanata* (30–32), and two from *A. thaliana*, AtSDR5 (AT2G47140) and AtSDR3 (AT2G47130) (10), have been shown to accept cardenolide intermediates *in vitro*, but *in planta* evidence for involvement in cardenolide biosynthesis only exists for Dl3βHSD1 (30) (Figures 6A, S15). Of the species included in the *KSI* gene tree, only *E. cheiranthoides* contained more than one copy (Figures 6B, S16). To our knowledge, no enzymes from this group have been biochemically characterized prior to this study.

**Figure 6.**
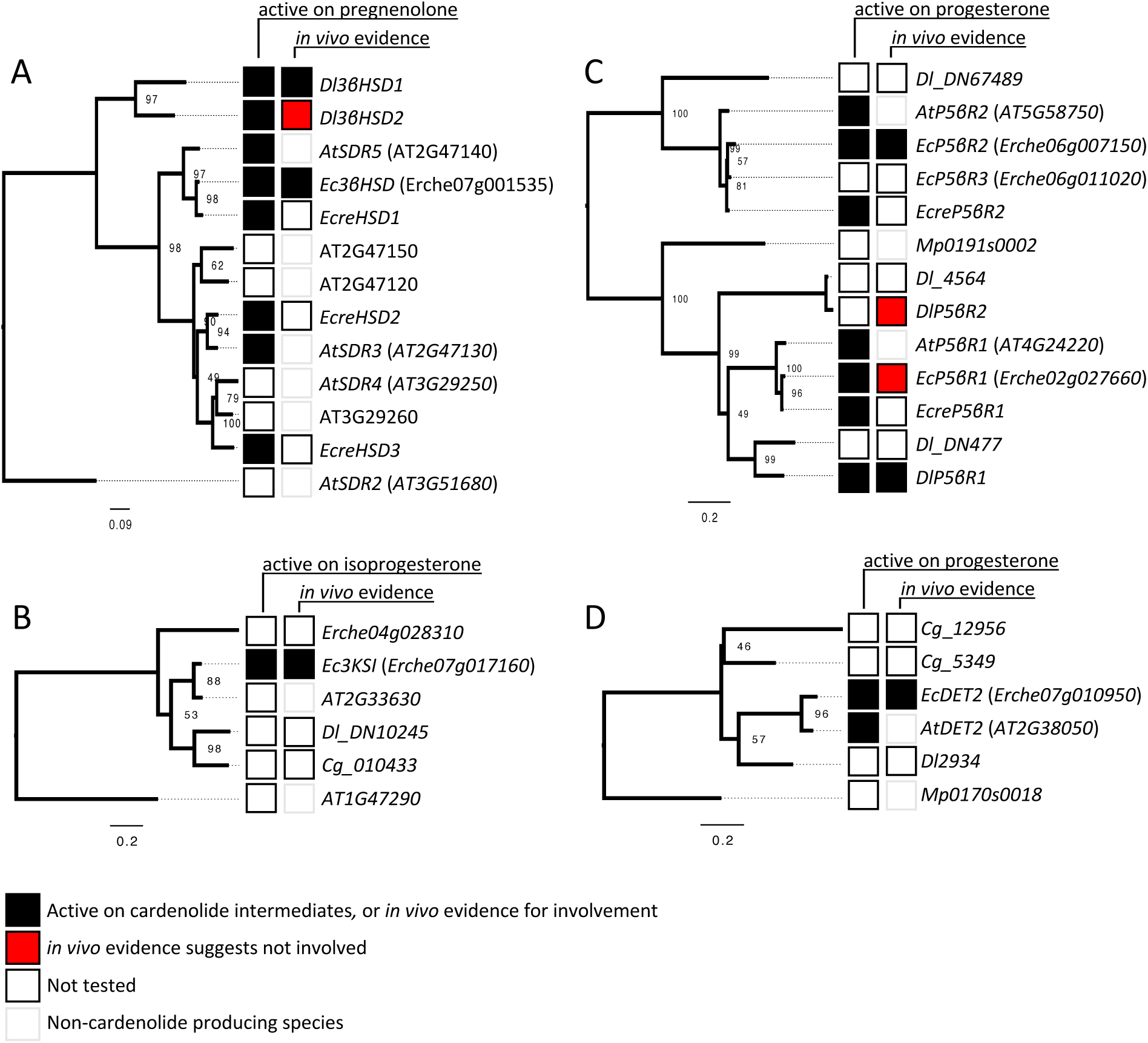
Phylogenetic analysis of *Erysimum cheiranthoides* cardenolide biosynthetic genes. Nucleotide phylogenies for cardenolide biosynthetic genes from *E. cheiranthoides* and selected other species for which functional work has been done. (A) 3β-hydroxysteroid dehydrogenase (3βHSD) and related short-chain dehydrogenases (SDR), (B) 3-ketosteroid isomerase (3KSI), (C) progesterone 5β-reductase (P5βR), and (D) steroid 5α-reductase (DET2). Experimental evidence for activity on cardenolide intermediates or *in vivo* evidence for involvement in cardenolide biosynthesis, either in this or previous studies, are marked by black squares. Enzymes for which *in vivo* evidence suggests the enzyme is not involved in cardenolide biosynthesis are marked by red squares. Species included: *Arabidopsis thaliana* (*At*/AT), *Calotropis gigantea* (*Cg*), *Erysimum cheiranthoides* (*Ec*/Erche), *Erysimum crepidifolium* (*Ecre*), *Digitalis lanata* (*Dl*), and *Marchantia polymorpha* (*Mp*). Numbers at nodes indicate bootstrap support from 10,000 replicates and scale bar indicates estimated substitutions per site. Multiple sequence alignments underlying these phylogenies are provided in Figures S15-S18.

*P5βR* genes fall into two clades arising from an ancient duplication event (33.8% amino acid identity between EcP5βR1 and EcP5βR2). Members of both clades have been shown to act on progesterone *in vitro* (13, 34, 45, 46), but only *DlP5βR1*, which is a more closely related to *EcP5βR1*, has a confirmed role in 5β-cardenolide biosynthesis (47). *EcP5βR2*, which we show to be required for 5β-cardenolide biosynthesis in *E. cheiranthoides*, belongs to the other clade (Figures 6C, S17). *EcDET2*, which is required for 5α-cardenolide biosynthesis in *E. cheiranthoides*, is a single copy gene for most species examined, and it accepts progesterone as a substrate even in species that do not make cardenolides, such as *A. thaliana* and *S. lycopersicum* (40). Intriguingly, *C. gigantea*, which accumulates 5α-cardenolides, contains two copies of this gene (Figures 6D, S18).

## DISCUSSION

### Identification E. cheiranthoides cardenolide biosynthesis genes

In this study, we identified and biochemically characterized four cardenolide biosynthetic enzymes from *E. cheiranthoides*. Of these, Ec3βHSD and EcP5βR2 belong to enzyme families that have been speculated, or shown, to be involved in cardenolide biosynthesis in other species (13, 30, 31, 45, 47). However, the mutant lines generated in this study provide critical *in vivo* evidence for their role in cardenolide synthesis, and the lack of a cardenolide phenotype for EcP5βR1 knockouts highlights the potential disparity between *in vitro* activity and *in vivo* function.

We additionally demonstrate the capacity of some enzymes to assume multiple roles in related metabolic pathways. The dwarf phenotype of the *det2* mutant lines in this study confirms that DET2 is required for brassinosteroid production in *E. cheiranthoides*. However, the lack of cardenolides with a fully saturated ring system in the *p5br2/det2* double mutants show that EcDET2 can also contribute to 5α-cardenolide synthesis. Because wildtype *E. cheiranthoides* primarily produces 5β-cardenolides, it is perhaps unsurprising that it does not have a dedicated copy of DET2 for cardenolide biosynthesis that could be regulated independently or have altered kinetics. In other *Erysimum* species that naturally accumulate higher levels of 5α-cardenolides, *DET2* may have been duplicated, although we were unable to confirm this with the available transcriptomes.

The involvement of a dedicated ketosteroid isomerase in cardenolide biosynthesis has been the subject of substantial research (38, 39), with others speculating that isomerization is catalyzed by 3βHSD, as is the case in animals (8, 48). Activity-guided fractionation of *Digitalis* protein extracts revealed that isomerization was catalyzed by an enzyme distinct from 3βHSD, but the protein sequence of the putative 3-ketosteroid isomerase was not identified. We identified Ec3KSI, a member of short chain dehydrogenase/reductase family 42E (SDR42E), as responsible for catalyzing the isomerization of isoprogesterone **3** to progesterone **4** (Figure 3) while lacking 3βHSD activity. Bacterial ketosteroid isomerases have been extensively characterized (49), but less is known about eukaryotic enzymes with KSI activity that lack 3βHSD activity. We demonstrated with CRISPR/Cas9-generated knockout lines that Ec3KSI is involved in cardenolide biosynthesis. However, even in the absence of Ec3KSI, 5β-cardenolides are produced, albeit in lower quantities than occur in wildtype plants. It is possible that some isomerization occurs non-enzymatically, or another enzyme, possibly Ec3βHSD, can partially compensate for the loss of KSI activity in the mutant lines.

Still in question is the extent to which plant 3βHSDs also possess KSI activity. While there has been at least one report of a tomato 3βHSD with KSI activity, our assays were inconclusive, as we saw isomerization when Ec3βHSD was supplied with pregnenolone **2** and NAD+, but not when Ec3βHSD is supplied with isoprogesterone **3**. This implies either that the isomerization by Ec3βHSD must accompany oxidase activity, the isomerization observed during the 3βHSD assay occurred non-enzymatically, or isomerization catalyzed by Ec3βHSD is slow. A comprehensive analysis of 3βHSD activity across plants, with careful attention to the spontaneous isomerization of isoprogesterone **3**, is warranted to differentiate these possibilities.

### Natural variation in P5βR expression and sequence influences cardenolide stereochemistry in Erysimum

We observed substantial interspecific variation in the accumulation of 5α-, 5β-, and dehydro-cardenolides across the *Erysimum* genus. Although cardenolides of each type had been reported previously in at least one *Erysimum* species (22, 24, 50), other studies focused primarily on 5β-cardenolides (45, 51). We found that the 5β-cardenolides digitoxigenin and cannogenin were restricted to a monophyletic clade containing *E. cheiranthoides*, *E. sylvestre*, and two closely related species of uncertain taxonomic identity (43), and that their occurrence coincided with expression of a functional *P5βR2-1*. In one monophyletic group of five *Erysimum* species, *P5βR2-1*, while expressed, contains a frameshift mutation, corresponding with the apparent loss of digitoxigenin and cannogenin synthesis. Several other species, including *E. crassipes*, *E. rhaeticum*, and *E. kotschyanum*, express *P5βR2-2* at high levels, but we did not detect digitoxigenin or cannogenin glycosides in these species. We hypothesize that the *P5βR2-1* clade is specialized for involvement in 5β-cardenolide biosynthesis, whereas *P5βR1* and *P5βR2-2* assume other roles in plant metabolism. The fact that these enzymes are expressed and active on cardenolide intermediates, but seem to be uninvolved in the pathway, points to the possibility of substrate channeling or compartmentalization of cardenolide biosynthesis.

By contrast, strophanthidin, a more hydroxylated 5β-cardenolide, was ubiquitous across the genus, and its occurrence did not depend on expression of *P5βR2-1* in most species. The occurrence of C5-hydroxylated 5β-cardenolides in the absence of *P5βR2-1* implies the existence of a P5βR-independent pathway for strophanthidin biosynthesis in *Erysimum*. For example, it is possible that inversion of carbon 5 stereochemistry occurs during C5-hydroxylation, and in fact a similar stereochemical inversion is thought to occur during C14-hydroxylation in cardenolide biosynthesis (52). Alternatively, we cannot exclude the possibility that one of the other expressed P5βR proteins forms digitoxigenin as an intermediate which is entirely converted by cardenolide hydroxylases to strophanthidin. Interestingly, the *E. cheiranthoides p5br2* mutant lacks strophanthidin, suggesting that such a P5βR-independent strophanthidin pathway, if it exists, may have been lost in this species.

Previous studies have reported digitoxigenin in several of the species for which we failed to detect it in our assays (22, 24). This discrepancy may be explained in a number of ways. First, we only sampled single accessions or seed batches per species, which may underestimate potentially substantial intraspecific variation in plant chemistry. Second, many early studies used paper chromatography or other low-resolution chromatographic techniques that would render correct stereochemical assignment difficult or impossible, and even high-resolution methods may miss differences in stereochemistry without appropriate reference material. For example, in a previous study profiling intact cardenolides in the same species studied here, the *E. cheiranthoides* clade exhibited a distinct chemotype with limited overlap in non-hydroxylated cardenolides compared to the other species examined, which may have been a reflection of a difference in carbon 5 stereochemistry (43). In particular, the *E. cheiranthoides* clade accumulated a set of unique mono- and diglycosides of digitoxigenin, cannogenol, and cannogenin, whereas most other *Erysimum* species accumulated an isomeric set of putative 5α-cardenolides with shifted HPLC retention times. Consistent with our results, no equivalent pattern was apparent for strophanthidin glycosides, and no 5α-isomers of strophanthidin have been described for any *Erysimum* species (22, 43).

Because core pathway enzymes are apparently active regardless of carbon 5 stereochemistry, switching between production of digitoxigenin and uzarigenin-glycosides only requires alteration to the expression or sequence of a single enzyme, P5βR2-1. Control of the production of unsaturated cardenolides, which do not occur at high levels in any of the species in this study, appears to be somewhat more cryptic, with potential for expression level, gene duplication, or protein-protein interactions to play a role. An analogous process controls steroidal glycoalkaloid diversity in *Solanum*, where expression of GAME25, a SDR related to Ec3βHSD, controls saturation of steroidal glycoalkaloids across the genus (53). Along with overall polarity, stereochemical configuration is a major contributor to variation in toxicity and deterrent activity of cardenolides (25–27, 54). As such, altering cardenolide stereochemical configuration may be a relatively simple evolutionary mechanism through which *Erysimum* fine-tunes its defensive profile to the most pervasive herbivores in a given ecological context. This theory is somewhat borne out by the findings of Mirzaei *et al*. (33), who showed that an *E. cheiranthoides P5βR2* mutant line, which produces primarily 5α-cardenolides, was more resistant to *Trichoplusia ni* (cabbage loopers) and *Myzus persicae* (green peach aphids), but extracts from 5β-cardenolide-producing wildtype plants had a greater inhibitory effect on porcine Na^+^,K^+^-ATPase *in vitro*.

This work represents a step forward in our understanding of the biosynthesis of medically important cardiac glycosides and provides insight into molecular mechanisms through which cardenolide structural variation may be controlled. The apparently modular nature of the cardenolide pathway, where the presence or activity of individual enzymes alters the pathway end products, has broad implications for future engineering of the pathway in heterologous systems for research or medical purposes. Such flexibility in pathway assembly may allow for rapid production and testing of varied cardenolide structures for biomedical applications.

Furthermore, the mutant *E. cheiranthoides* lines presented here will facilitate investigation of the functional and ecological implications of carbon 5 configuration in cardenolides. Insect feeding assays and field experiments with these mutant plants would illuminate herbivore preferences and provide further insight into the selective pressures that may have shaped the cardenolide profiles observed in nature today.

## MATERIALS AND METHODS

### Plant growth, cloning, expression, and knockout of candidate genes

Plant growth, cloning of candidate genes, transient expression in *Nicotiana benthamiana*, and CRISPR/Cas9 knockout in *Erysimum cheiranthoides* were performed as described previously (35). For the genus-wide experiment, lyophilized tissue collected during a previous study and stored at -20 °C (43) was used. Primers used for cloning of candidate genes and for generation and screening of CRISPR/Cas9 mutants are provided in Table S9.

### Mutagenic screens

Ethyl methanesulfonate (EMS) mutagenesis was modified from Mirzaei et al., 2020 (33). Ten grams of *E. cheiranthoides* seeds were soaked at 4 °C overnight in 100 mL of 100 mM phosphate buffer, pH 7.5. The buffer was decanted, and the seeds were resuspended in 100 mL of fresh phosphate buffer with 0.6% EMS (Sigma-Aldrich, St. Louis, MO). The seeds were shaken at 23 °C for six hours, washed twenty times with deionized water, and grown in twenty 25×25 cm flats to maturity. M2 seeds were pool-harvested from each flat at two timepoints. To screen for mutants in cardenolide biosynthesis, 32 plants from each pool (in total, 1120 plants) were grown for four weeks. Approximately 30 mg of leaf tissue was harvested from each plant for UHPLC-MS analysis. Plants that showed divergent cardenolide phenotypes were backcrossed to wildtype, and F2 progeny were used for bulked segregant analysis (BSA) as described previously (33).

### Coexpression networking analysis

Raw RNA-sequencing reads from 48 *Erysimum* species (43) were downloaded from the NCBI Short Read Archive (SRP225657) and were pseudoaligned to the transcriptome associated with *E. cheiranthoides* genome v2.1 (NCBI: PRJNA563696)(55, 56) using kallisto (57) with default parameters, yielding transcript counts, which were filtered to retain transcripts with more than 10 counts in at least 10 samples. Filtered counts were used for the mr2mods gene coexpression analysis pipeline using default parameters (58).

### Protein expression and purification

For protein purification, genes were inserted into the Champion^TM^ pET300-NT-DEST plasmid (ThermoFisher Scientific, Waltham, MA), before being transformed into Rosetta(DE3) *E. coli* (MilliporeSigma, St. Louis, MO). Single colonies were picked from LB agar plates (100 μg/mL carbenicillin; 20 μg/mL chloramphenicol) to inoculate 10 mL liquid LB cultures with the same antibiotics. Cultures were grown overnight at 37 °C and 225 rpm in a I2500 Incubator Shaker incubator (Eppendorf, Hamburg Germany). After 18 hours, 6 mL of the culture was transferred to 250 mL TB medium (59) containing the same antibiotics, and cells were grown under the same conditions until OD_600_=0.6. Isopropyl β-d-1-thiogalactopyranoside (IPTG, ThermoFisher Scientific) was added to a final concentration of 1 mM to induce protein expression, and cultures were incubated at 25 °C and 225 RPM for 6 hours before being placed on ice and centrifuged at 5,000 rcf and 4 °C for 15 minutes in a Sorvall RC5C Plus centrifuge (ThermoFisher Scientific). Cells were resuspended in 40 mL of lysis buffer, consisting of 40 mM Tris pH8, 20 mM imidazole (Sigma-Aldrich, St. Louis, MO), 500 mM NaCl (ThermoFisher Scientific), 10% (v/v) glycerol, and 1% (v/v) Tween 20 (Sigma-Aldrich). The cell suspension was lysed by freezing in liquid N_2_ and thawing on ice twice, followed by sonication using a Branson Sonifier 250 while still on ice, four times in 10-second intervals, with a 30-second rest between each interval. The cell lysate was centrifuged for 45 minutes at 4 °C at 13,000 rcf. Following centrifugation, the supernatant was loaded onto a column containing 1 mL of Ni-NTA resin (Invitrogen, Waltham, MA) that had been previously equilibrated with 1 column volume of lysis buffer. After all supernatant had passed through the column, 1 column volume of wash buffer (50 mM Tris pH 8.0, 20 mM imidazole, 500 mM NaCl, 10% (v/v) glycerol) was passed through the column.

Finally, 0.5 mL elution buffer (50 mM Tris pH 8.0, 250 mM imidazole, 500 mM NaCl, 10% v/v glycerol) was loaded onto the column, and the flow-through was collected in a 2 mL microcentrifuge tube (Laboratory Products Sales, Rochester, NY). Protein concentration was measured using a NanoDrop One (ThermoFisher Scientific).

### In vitro enzyme assays

Steroid-3β-hydroxysteroid dehydrogenase (3βHSD) and 3-ketosteroid isomerase (3KSI) assay conditions were adapted from previous studies (39, 53). One hundred μl reactions contained 4 mM KPO_4_ buffered at pH 6.5 and 1 μg purified enzyme. For the 3βHSD oxidation, final concentrations of 150 μM NAD+ (Sigma-Aldrich) and 10 μM pregnenolone (Sigma-Aldrich) were used. To test for 3βHSD reductase activity, 150 μM NADH (Sigma-Aldrich) and 10 μM 5β-pregnane-3,20-dione (aablocks, San Diego, CA) or 5α-pregnane-3,20-dione (Sigma-Aldrich) were used. For the KSI assay, the same conditions were used, but 10 μM isoprogesterone (TLC Pharmaceutical Standards, Newmarket, ON) was used and NAD+/NADH were omitted.

Progesterone 5β-reductase (P5βR) assays were adapted from Herl et al. (13) and Sonawane et al. (53). 100 μl reactions contained final concentrations of 4 mM KPO_4_ pH 7.2, 150 μM NADPH (Cayman Chemical, Ann Arbor, MI, USA), 10 μM progesterone (Sigma-Aldrich), and 1 μg purified enzyme.

Reactions were incubated for 1 hour at 28 °C for the 3βHSD and 3KSI assays, and at 37°C for the P5βR assay. All assays were terminated by addition of 100 μl 100% methanol (ThermoFisher Scientific) containing 15 μg/mL ouabain (Sigma-Aldrich) as an internal standard, centrifuged at 17,000 rcf in an Eppendorf 5417R Centrifuge at 4 °C, and transferred to vials (ThermoFisher Scientific) for UHPLC-MS analysis.

### Metabolite extraction

Metabolites were extracted from fresh tissue of *E. cheiranthoides* and *N. benthamiana* as described previously (35). Where indicated, metabolite extracts were subjected to acid hydrolysis to isolate cardenolide genins using a protocol adapted from Schaller & Kries (19). In brief, 700 μL 100% methanol for two leaf 14 mm leaf disks of fresh *E. cheiranthoides* tissue or 750 μL 95% (v/v) methanol per 20 mg of lyophilized tissue in the genus-wide experiment, was used to extract metabolites for 30 minutes at 25 °C. After centrifugation for three minutes at 17,000 rcf, 700 μL supernatant was transferred to a fresh microcentrifuge tube. Twenty μL of 6 M hydrochloric acid was added to each tube, and samples were incubated for 18 hours at 28 °C. Hydrolysis was terminated with 200 μL saturated sodium phosphate solution, and samples were extracted twice with 200 μL chloroform. The organic (lower) phase was evaporated to dryness in a Savant SpeedVac^TM^ SC110 (Thermo Fisher Scientific). Samples were resuspended in 50 μL methanol and centrifuged for 10 minutes at 17,000 rcf before being transferred to glass mass spectrometry vials for UHPLC-MS analysis.

### Liquid chromatography-mass spectrometry (LC-MS) analysis

All samples were analyzed on an UltiMate 3000 UHPLC system coupled to a Q-Exactive hybrid quadrupole-orbitrap mass spectrometer (Thermo Fisher Scientific, Waltham, MA). The instrument was fitted with a Supelco Titan^TM^ C18 UHPLC Column (80Å, 100 x 2.1 mm, particle size 1.9 μm; Sigma Aldrich). Injections of 2 μL were separated by a solvent gradient consisting of mobile phase A (water + 0.1% (v/v) formic acid) and mobile phase B (acetonitrile + 0.1% (v/v) formic acid). A 13-minute method was used for analysis of non-hydrolyzed samples: 0-0.55 minutes, hold at 2% B; 0.5-10 minutes, linear gradient from 2%-97% B; 10-11.5 minutes, hold at 97% B, 11.5-13 minutes, hold at 2% B. A longer solvent gradient was used in hydrolysis experiments: 0-5 minutes, hold at 2% B; 5-22 minutes, linear gradient from 2%-97% B; 22-23.5 minutes, hold at 97% B, 23.5-25 minutes, hold at 2% B. All solvents were Optima LC/MS grade (Thermo Fisher Scientific). The solvent flow rate was 0.5 mL/minute, the column oven was set to 40 °C, and the autosampler temperature was 15 °C for all methods. The mass spectrometer was run in full scan positive ionization mode. Targeted MSMS spectra were collected with an isolation window of 2.0 *m/z* and normalized collision energy of 30%.

LC-MS peak areas were quantified using a custom processing method in Xcalibur^TM^ Software (ThermoFisher Scientific) using the following parameters: peak detection ICIS, smoothing points 1, baseline window 40, area noise factor 5, peak noise factor 15, tailing factor 2. Mass features used for quantification are provided in Table S10 for in-tact cardenolides, Table S11 for hydrolyzed cardenolides, and Table S12 for cardenolide intermediates from *in vitro* and *N. benthamiana* assays.

### Statistical and phylogenetic analysis

The following functions in R statistical software (60) were used for statistical tests, which were performed on log-transformed LC-MS peak areas, normalized to an internal standard: aov, TukeyHSD, and t.test. Plots were made using MSnbase (61, 62), multcompView (63), and pheatmap (64).

Sequences homologous to *Ec3βHSD*, *EcKSI*, *EcP5βR2*, and *EcDET2* were identified using BLAST against publicly available transcriptomes for *Arabidopsis thaliana* (65), *Calotropis gigantea* (66), *Digitalis lanata* (48) (NCBI PRJNA923725), *Marchantia polymorpha* (67), and other *Erysimum* species (43) (NCBI PRJNA563696), and were aligned using ClustalW (68, 69). Gene phylogenies were inferred using IQ-TREE web server (70–72) with default parameters, except bootstrap alignments were increased to 10,000. Raw data underlying all figures are available in the Supporting Information.

## ACKNOWLEDGMENTS

We thank Tobias Krug for assistance with laboratory assays. This research was funded by United States Department of Agriculture award 2020-67013-30896, US National Science Foundation award 1645256, and an award from the Triad Foundation to GJ; a Chemistry Biology Interface Training Program fellowship under National Institutes of Health/National Institute of General Medical Sciences (T32GM138826) and a US National Science Foundation Graduate Research Fellowship (DGE–2139899) to GCY; a Swiss National Science Foundation grant (PCEFP3-194590) to TZ; and a Summer Undergraduate Research Fellowship from the American Society of Plant Biologists and a Rawlings Cornell Presidential Research Scholar award to MLA.

## COMPETING INTERESTS

None declared.

## AUTHOR CONTRIBUTIONS

GCY, MLA, and GJ designed the research; GCY, MLA, and TZ performed the research; TZ contributed critical plant material; GCY and MLA analyzed data; GCY, TZ, and GJ wrote and edited the manuscript.

## DATA AVAILABILITY

The raw data that support the findings of this study are available in the Supporting Information. Seeds from mutant lines will be made available from the Arabidopsis Biological Resource Center. Due to high mortality and poor seed set, *DET2* knockout lines are not available from the ABRC.

## Supporting Information

The following Supporting Information is available for this article:

**Figure S1.**
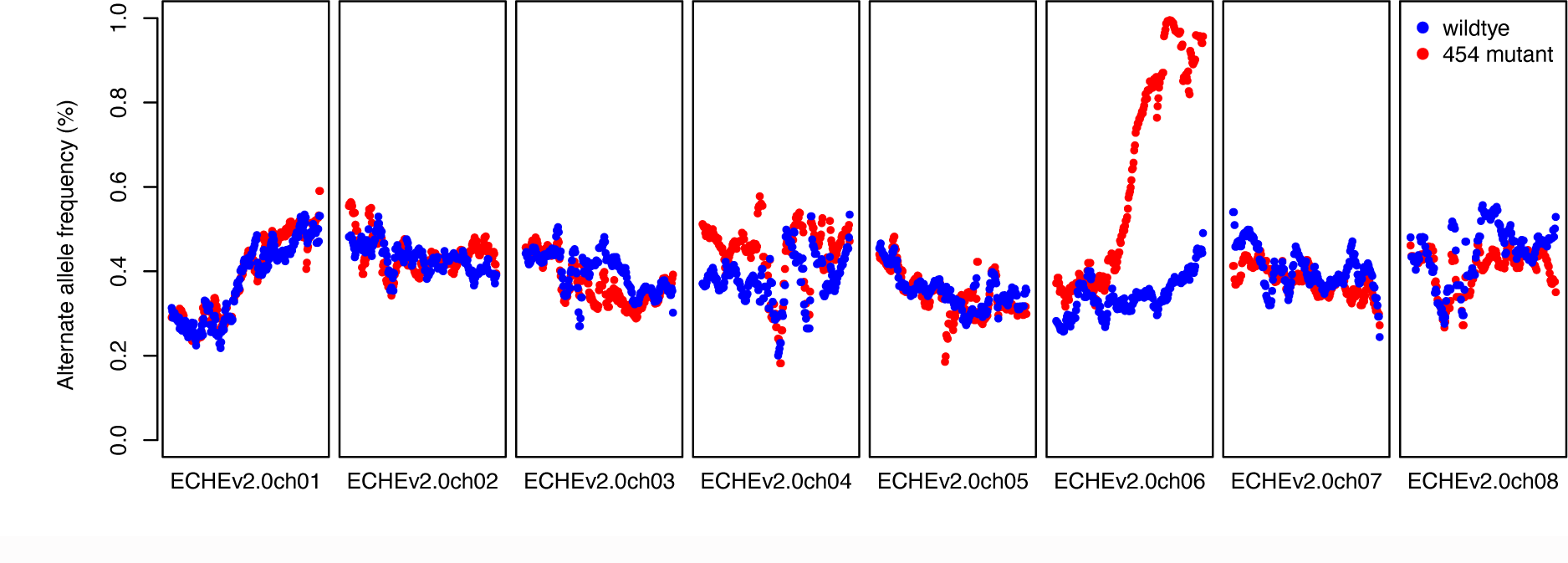
Bulked segregant analysis (BSA) from *Erysimum cheiranthoides* mutant #454. Mutant #454 was generated via ethyl methanesulfonate (EMS) mutagenesis. Alternate (mutant) allele frequency, smoothed over 1 Mbp segments, is plotted across the eight *E. cheiranthoides* chromosomes. In plants with a mutant chemotype (red), mutant alleles dominate in the latter half of chromosome seven. Within this region is a progesterone reductase (*EcP5βR2*) with two missense mutations (R184K and G201R) in mutant #454 plants. The cardenolide phenotype associated with mutant #454 and the BSA results displayed here were first described in Mirzaei et al. 2020. The causal mutation at the linked locus was first described in this study.

**Figure S2.**
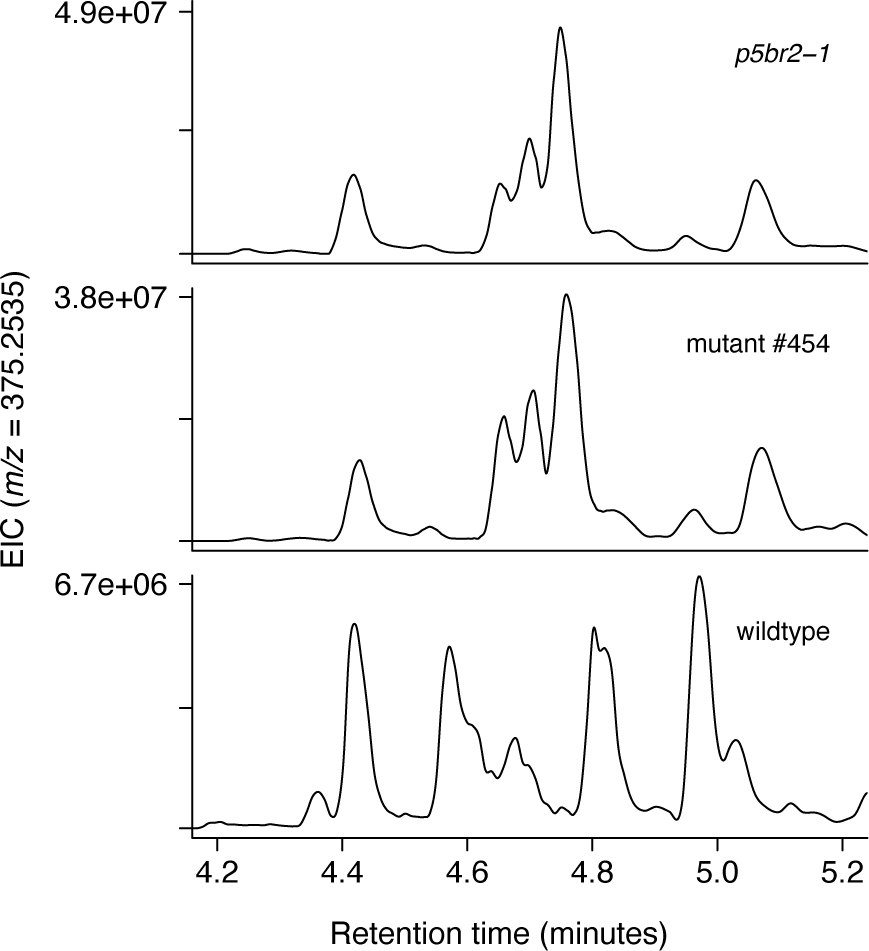
Extracted ion chromatograms (EIC) for cardenolides in wildtype *Erysimum cheiranthoides*, mutant #454, and *p5br2-1*. EIC for *m/z* 375.2535, a fragment common to digitoxigenin and uzarigenin glycosides, resulting from the neutral loss of all sugar moieties, leaving only the genin intact. In mutant #454 and *p5br2* mutant lines, we observe cardenolides with the same mass as those found in wildtype plants, but they elute at different retention times, suggesting that they may be structural isomers.

**Figure S3.**
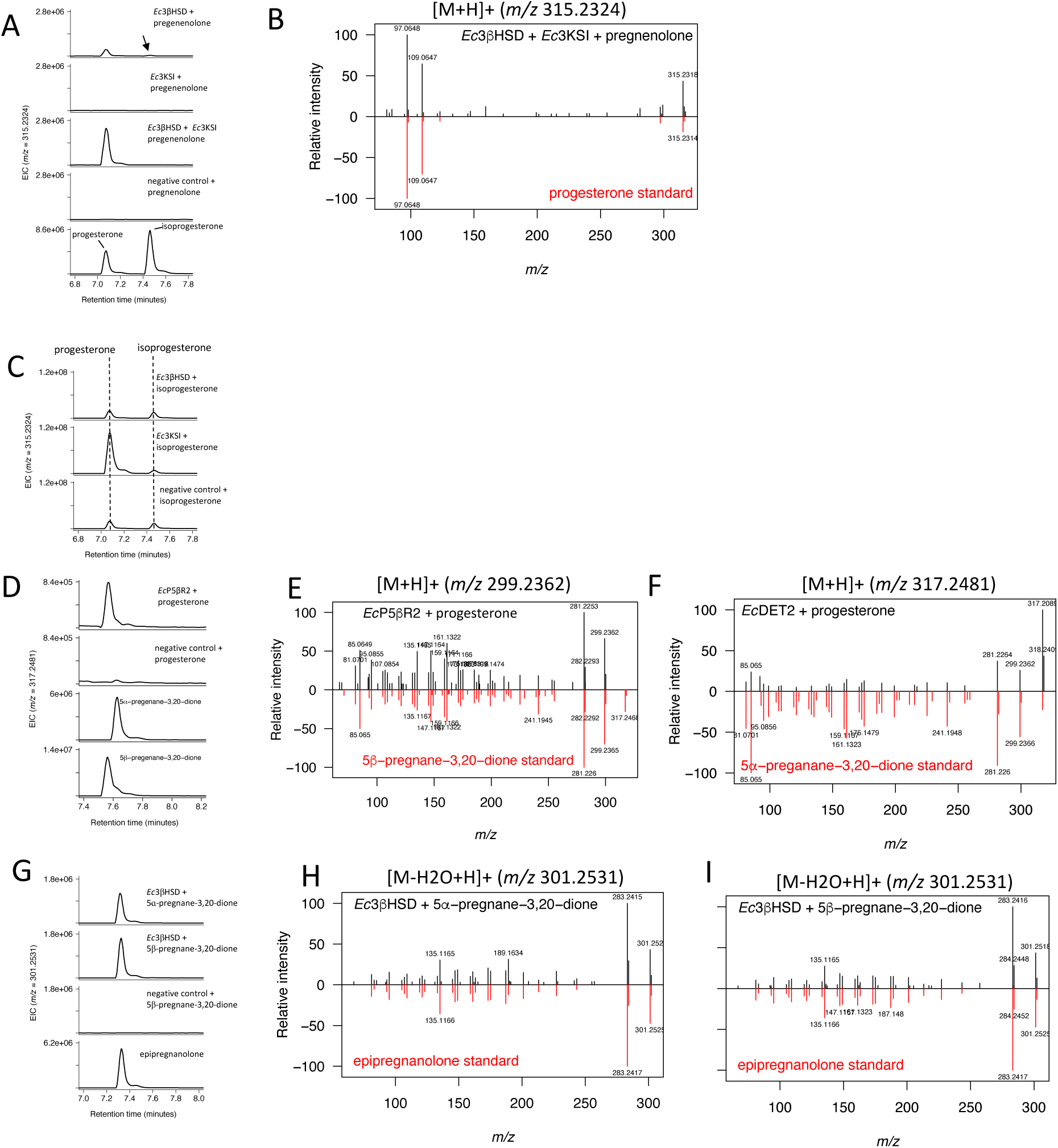
Extracted ion chromatograms (EIC) and MSMS spectra from products of *in vitro* or *Nicotiana benthamiana* transient expression assays compared with authentic standards. EIC (A) of isoprogesterone and progesterone formed by supplying *Ec*3βHSD or/and *Ec*3KSI with pregnenolone and NAD+ *in vitro*. (B) MSMS spectra of progesterone. High-quality isoprogesterone MSMS spectrum could not be collected from *in vitro* assays due to low signal. (C) EIC of conversion of isoprogesterone to progesterone *in vitro*. (D) EIC of 5β-pregnane-3,20-dione formed by supplying *Ec*P5βR2 with progesterone and NADPH *in vitro*, and corresponding MSMS spectrum (E). (F) MSMS spectrum of 5ɑ-pregnane-3,20-dione formed by *Ec*DET2 following coinfiltration with progesterone in leaves of *Nicotiana benthamiana*. EIC (G) and MSMS spectra of epipregnanolone or diastereomer formed by supplying *Ec*3βHSD with 5ɑ-pregnane-3,20-dione (H) or 5β-pregnane-3,20-dione (I) and NADH *in vitro*.

**Figure S4.**
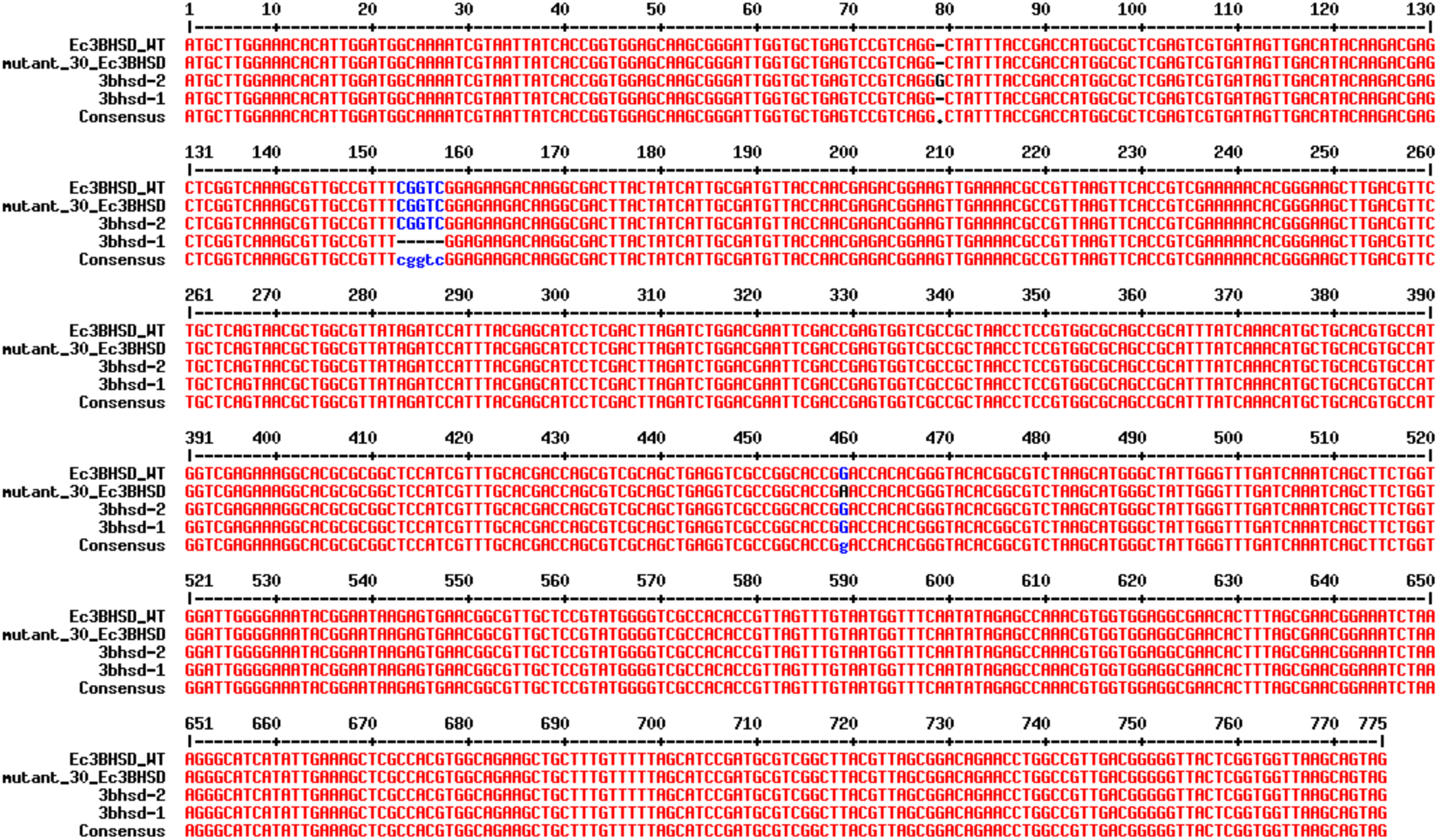
Aligned nucleotide coding sequences of Erysimum cheiranthoides 3βHSD from WT and mutant lines. Mutant #30 was generated via chemical mutagenesis with ethyl methanesulfonate (EMS), and *3bhsd-1* and *3bhsd-2* lines were generated with CRISPR/Cas9. Abbreviations: *Erysimum cheiranthoides* (Ec), wildtype (WT), 3β-hydroxysteroid dehydrogenase (3βHSD). Sequences of gRNAs used for generation of these lines are available in Table S1. MultAlin (http://multalin.toulouse.inra.fr/multalin/) was used to produce the alignment.

**Figure S5.**
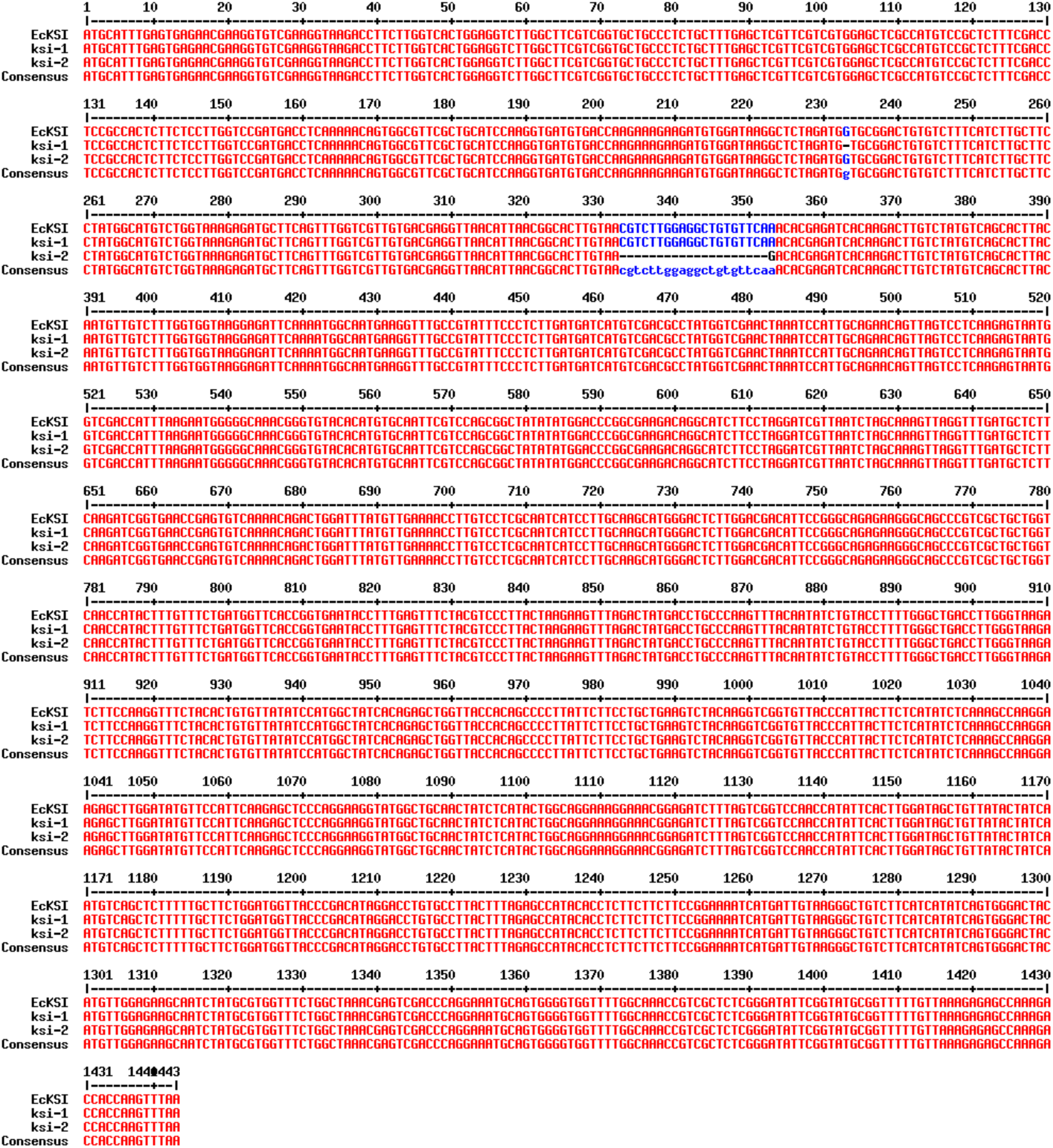
Aligned nucleotide coding sequences of Erysimum cheiranthoides 3KSI from WT and mutant lines. *3ksi-1* and *3ksi-2* lines were generated with CRISPR/Cas9. Abbreviations: *Erysimum cheiranthoides* (Ec), wildtype (WT), 3ketosteroid isomerase (3KSI). Sequences of gRNAs used for generation of these lines are available in Table S1. MultAlin (http://multalin.toulouse.inra.fr/multalin/) was used to produce the alignment.

**Figure S7.**
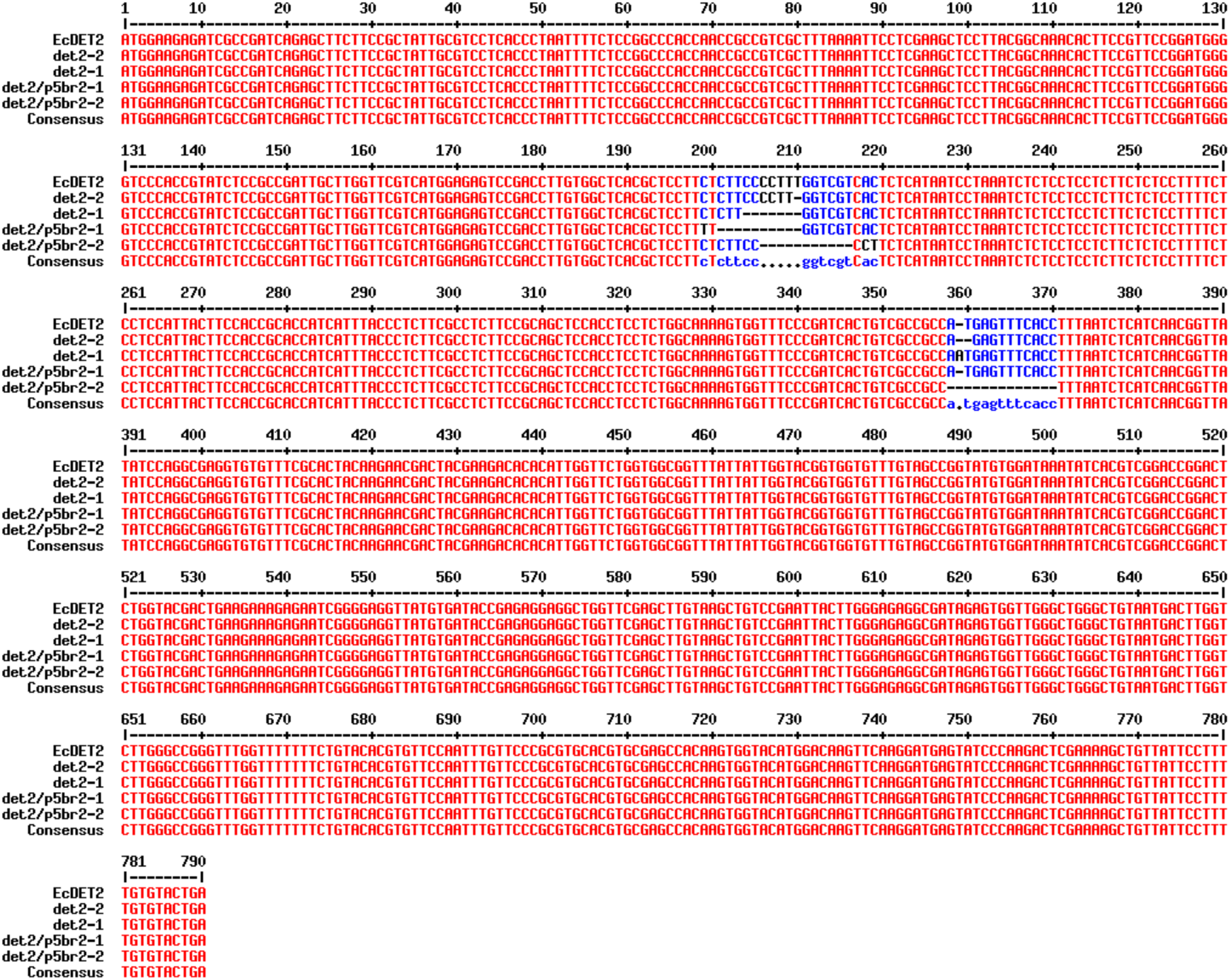
Aligned nucleotide coding sequences of Erysimum cheiranthoides DET2 from WT and mutant lines. *det2-1* and *det2-2* lines were generated with CRISPR/Cas9 in the WT background. *det2/p5br2-1* and *det2/p5br2-2* lines were generated with CRISPR/Cas9 in the *p5br2-1* background. Abbreviations: *Erysimum cheiranthoides* (Ec), wildtype (WT), progesterone 5β-reductase (P5βR), steroid 5α-reductase (DET2). Sequences of gRNAs used for generation of these lines are available in Table S1. MultAlin (http://multalin.toulouse.inra.fr/multalin/) was used to produce the alignment.

**Figure S8.**
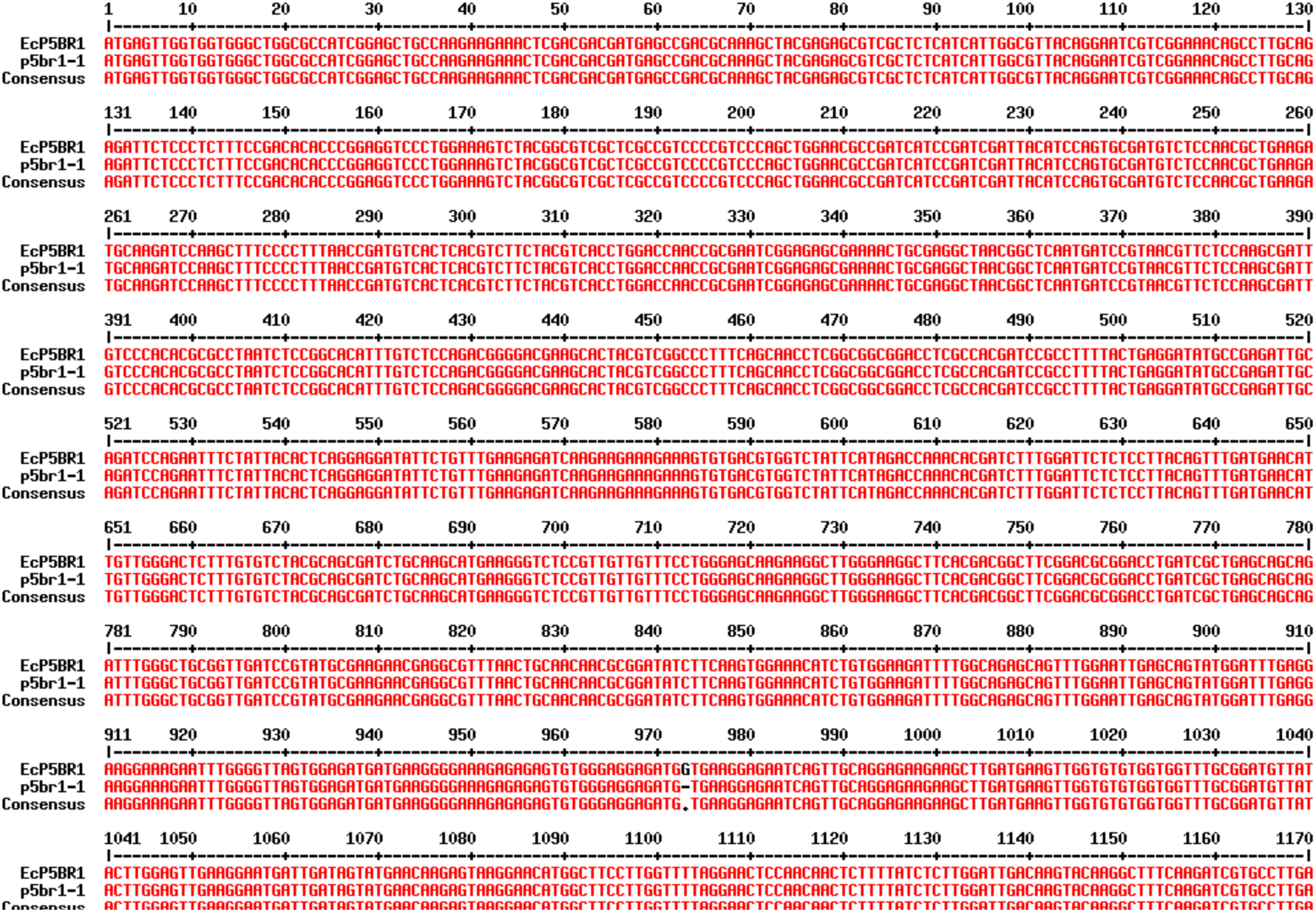
Aligned nucleotide coding sequences of Erysimum cheiranthoides P5βR1 from WT and mutant lines. The *p5br1-1* line was generated with CRISPR/Cas9. Abbreviations: *Erysimum cheiranthoides* (Ec), wildtype (WT), progesterone 5β-reductase (P5βR). Sequences of gRNAs used for generation of these lines are available in Table S1. MultAlin (http://multalin.toulouse.inra.fr/multalin/) was used to produce the alignment.

**Figure S9.**
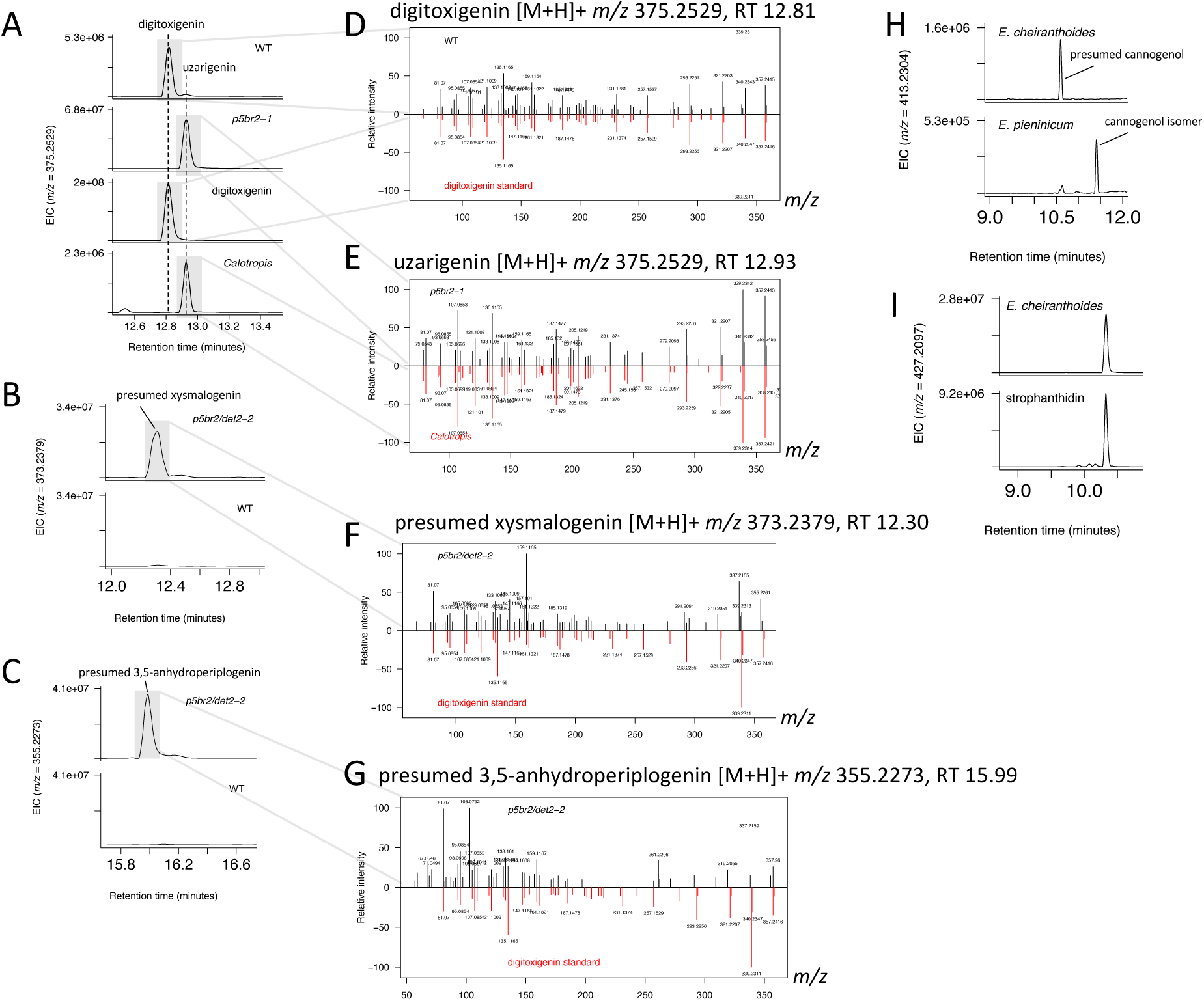
Extracted ion chromatograms and MSMS spectra for cardenolide genins. Selected extraction ion chromatograms (EIC) and MSMS spectra for cardenolide genins from hydrolyzed extracts of *Erysimum cheiranthoides* wildtype (WT) and mutant leaves, compared to authentic standards where available. (A) EIC of digitoxigenin and uzarigenin [M+H]+, which are stereoisomers and are separated by retention time. An uzarigenin standard was not available; hydrolyzed *Calotropis procera* leaf extract, which are known to contain uzarigenin, was used instead. (B) Presumed xysmalogenin [M+H]+ is abundant in *p5br2/det2* double mutants compared to WT leaves. 3,5-anhydroperiplogenin [M+H]+ (C) is formed when canarigenin glycosides are subjected to acidic conditions in *p5br2/det2* double mutants. (D-G) Corresponding MSMS spectra. Authentic standards were not available for xysmalogenin (F) or 3,5-anhydroperiplogenin (G). Instead, they are compared to a digitoxigenin MSMS spectrum. The spectra are similar, but some peaks are shifted by 2 Daltons, providing further evidence that these peaks represent dehydrocardenolides. (H) Cannogenol [M+Na]+ in *E. cheiranthoides*, and an isomer in *E. pieninicum* that may be the 5α conformation. (I) Strophanthidin [M+Na]+ in *E. cheiranthoides* compared to an authentic standard.

**Figure S10.**
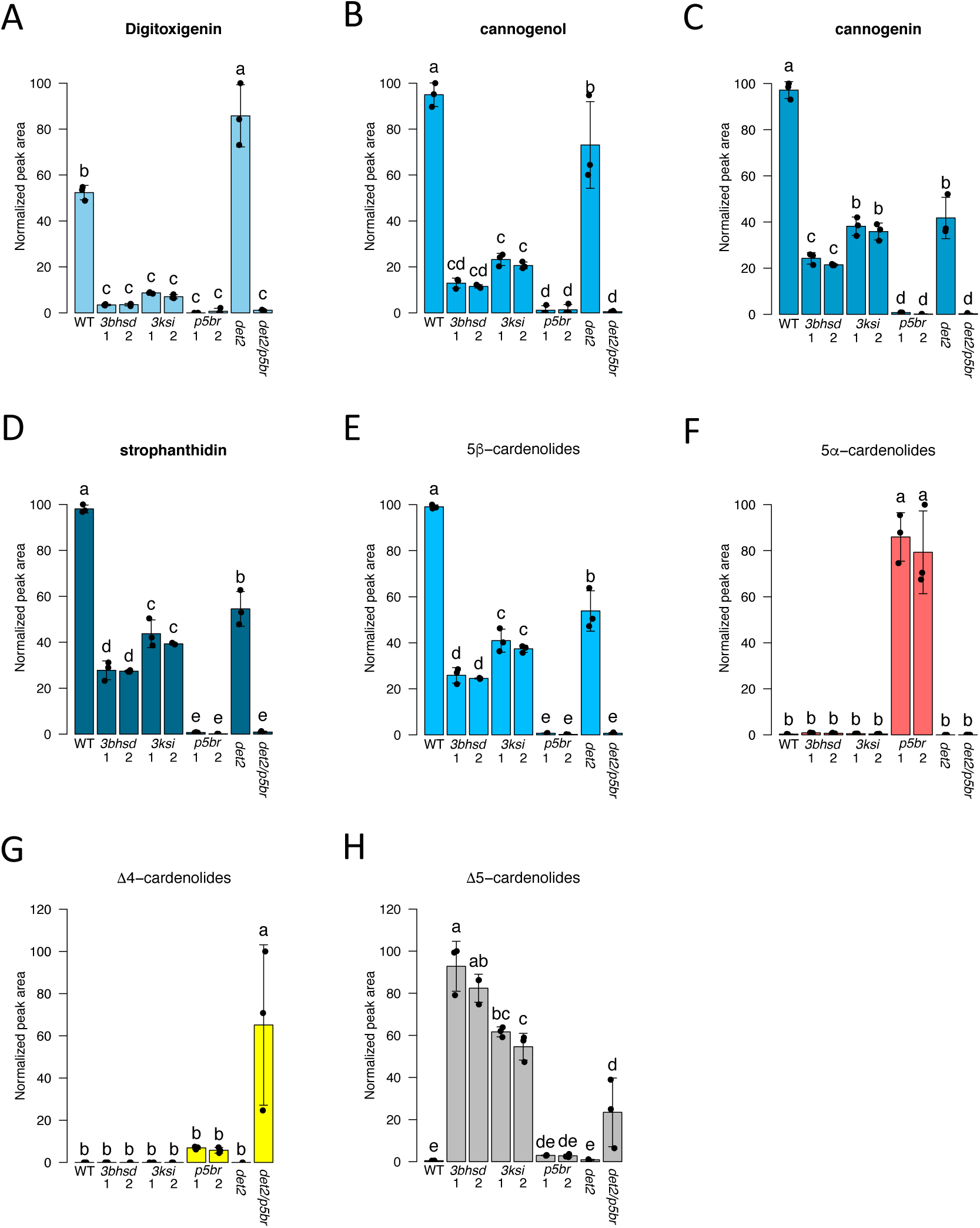
Normalized peak areas for cardenolide genins from hydrolyzed leaf extracts of *Erysimum cheiranthoides* mutant lines. (A) Digitoxigenin, (B) cannogenol, (C) cannogenin, (D) strophanthidin, (E) total 5β-cardenolides (sum of digitoxigenin, cannogenol, cannogenin, and strophanthidin), (F) 5α-cardenolides (uzarigenin), (G) Δ4-cardenolides (dianhydroperiplogenin as a proxy for canirigenin), (H) Δ5-cardenolides (xysmalogenin). Abbreviations: wildtype (WT), 3β-hydroxysteroid dehydrogenase (*3bhsd*), 3-ketosteroid isomerase (*3ksi*), progesterone 5β-reductase (*p5br*), and steroid 5α-reductase (*det2*). Error bars are ± s.d. Letters indicate P<0.05, one-way ANOVA with post-hoc Tukey’s HSD test.

**Figure S11.**
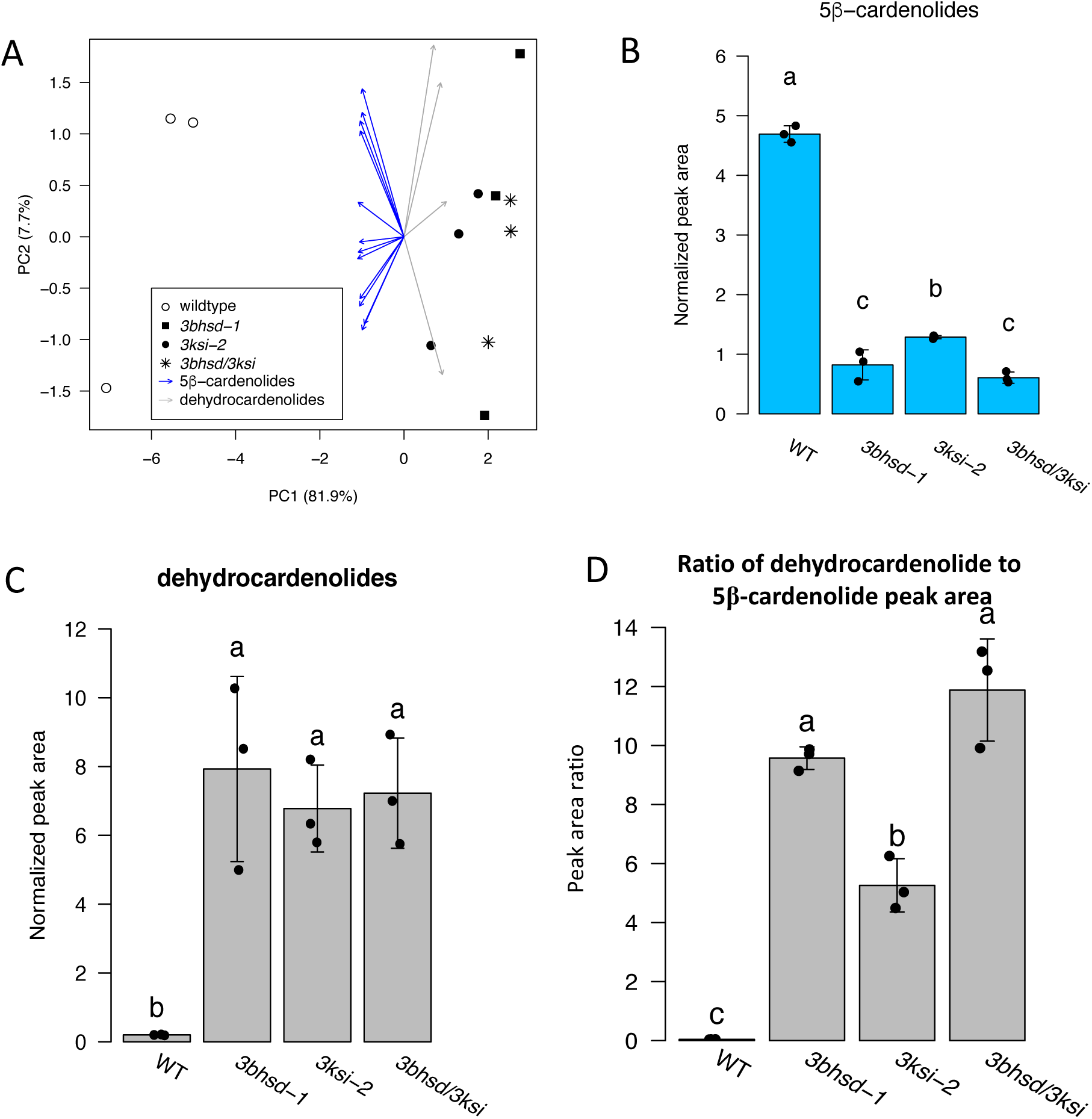
Cardenolide analysis from *Ec3βHSD*/*Ec3KSI* double mutants. (A) PCA of cardenolides detected in *Erysimum cheiranthoides* wildtype, *3bhsd*, *3ksi*, and *3bhsd/3ksi* mutant lines. Normalized peak area of (B) total 5β-cardenolides and (C) total dehydrocardenolides. (D) Ratio of dehydrocardenolide peak area to 5β-cardenolide peak area. Abbreviations: wildtype (WT), 3β-hydroxysteroid dehydrogenase (*3bhsd*), 3-ketosteroid isomerase (*3ksi*). N=3 plants per line. Error bars are ± s.d. Letters indicate P<0.05, one-way ANOVA with post-hoc Tukey’s HSD test.

**Figure S12.**
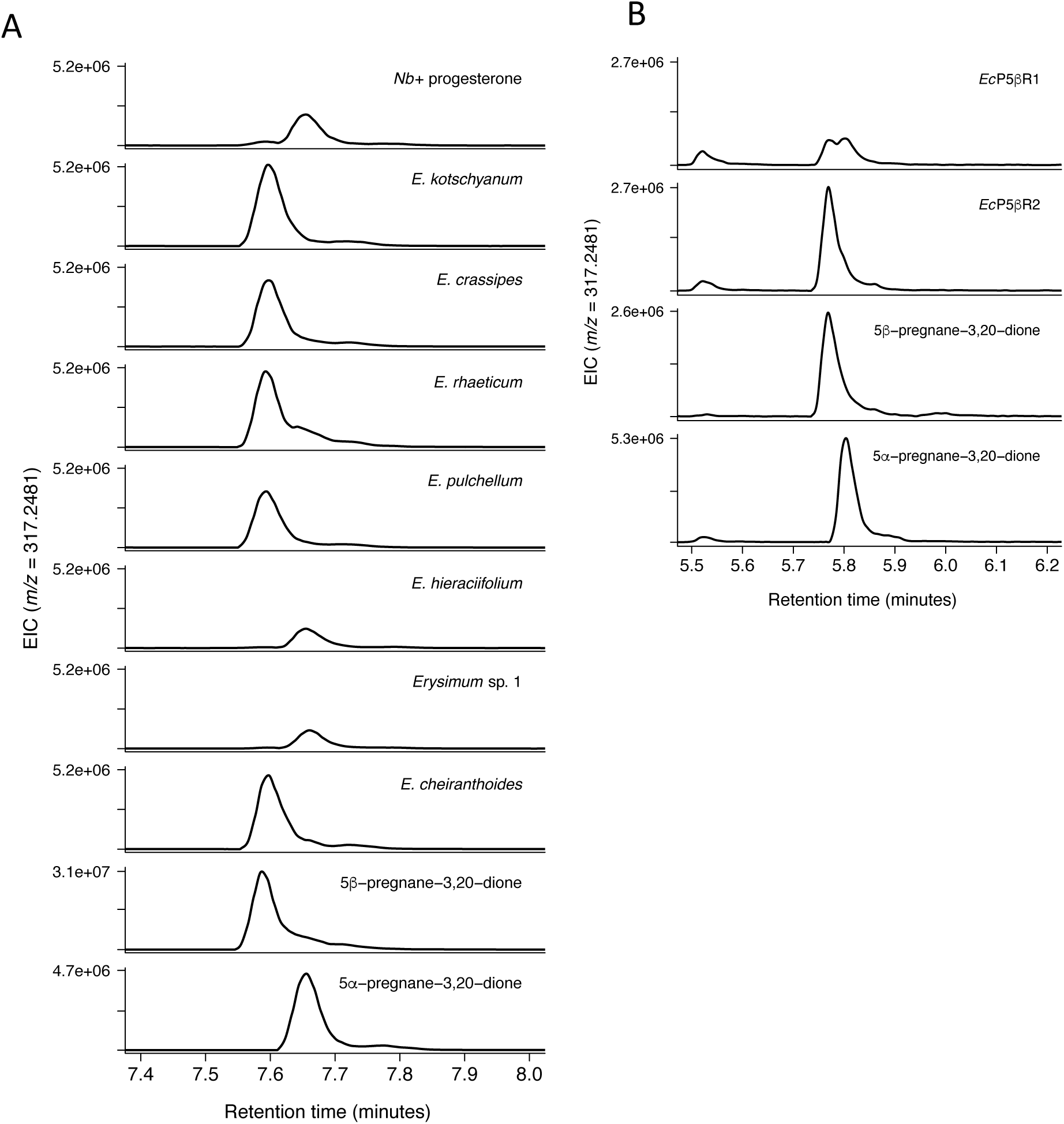
Coinfiltration of *Erysimum* progesterone 5β-reductases and progesterone in *Nicotiana benthamiana* leaves. *Erysimum* progesterone 5β-reductases (P5βR) cloned into pEAQ-HT-DEST1 were infiltrated into leaves of *N. benthamiana* (*Nb*), followed by infiltration of progesterone after three days. Extracted ion chromatograms (EIC) at *m/z*=317.2481 show production of 5β-pregnane-3,20-dione. When progesterone is infiltrated into *N. benthamiana* leaves with no co-infiltrated enzyme, 5α-pregnane-3,20-dione is produced by endogenous *N. benthamiana* enzymes. (A) P5βR2 orthologs from selected species of *Erysimum. E. hieraciifolium* and *Erysimum* sp. 1 P5βR2 proteins are truncated by a premature stop codon and are non-functional in this assay. (B) P5βR1 and P5βR2 from *E. cheiranthoides* (*Ec*). The retention time disparity between the two experiments is due to a shorter LCMS method used in panel B.

**Figure S13.**
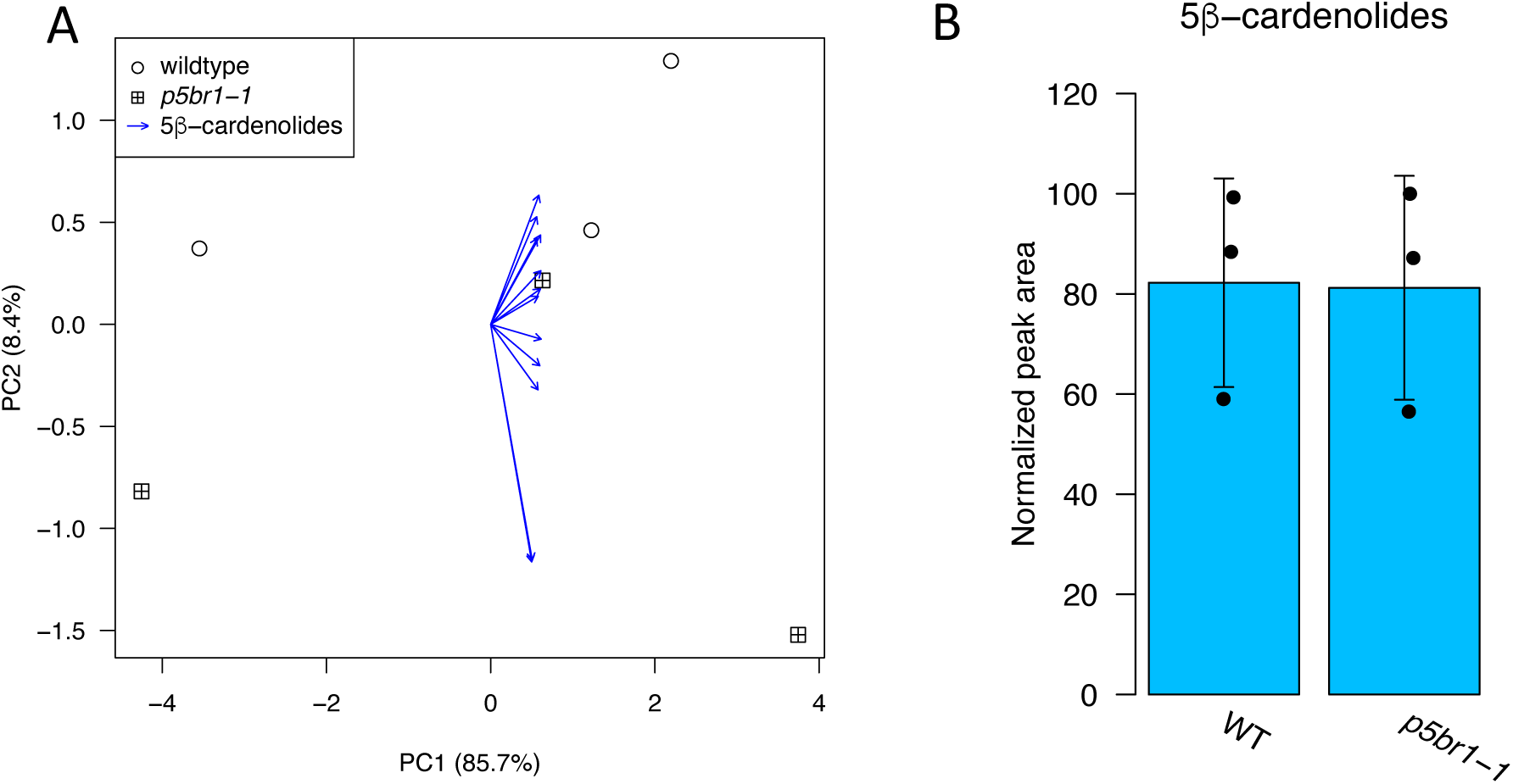
Cardenolide analysis from *EcP5βR1* mutants. (A) PCA of cardenolides detected in *Erysimum cheiranthoides* wildtype (WT) and *p5br1* (progesterone 5β-reductase 1) mutant line. (B) Normalized peak area of total cardenolides. All cardenolides in this experiment are also found in WT *E. cheiranthoides* and are presumed to be 5β-cardenolides. No dehydrocardenolides were detected. No differences were detected between groups (one way ANOVA: P = 0.957). N=3. Error bars are ± s.d.

**Figure S14.**
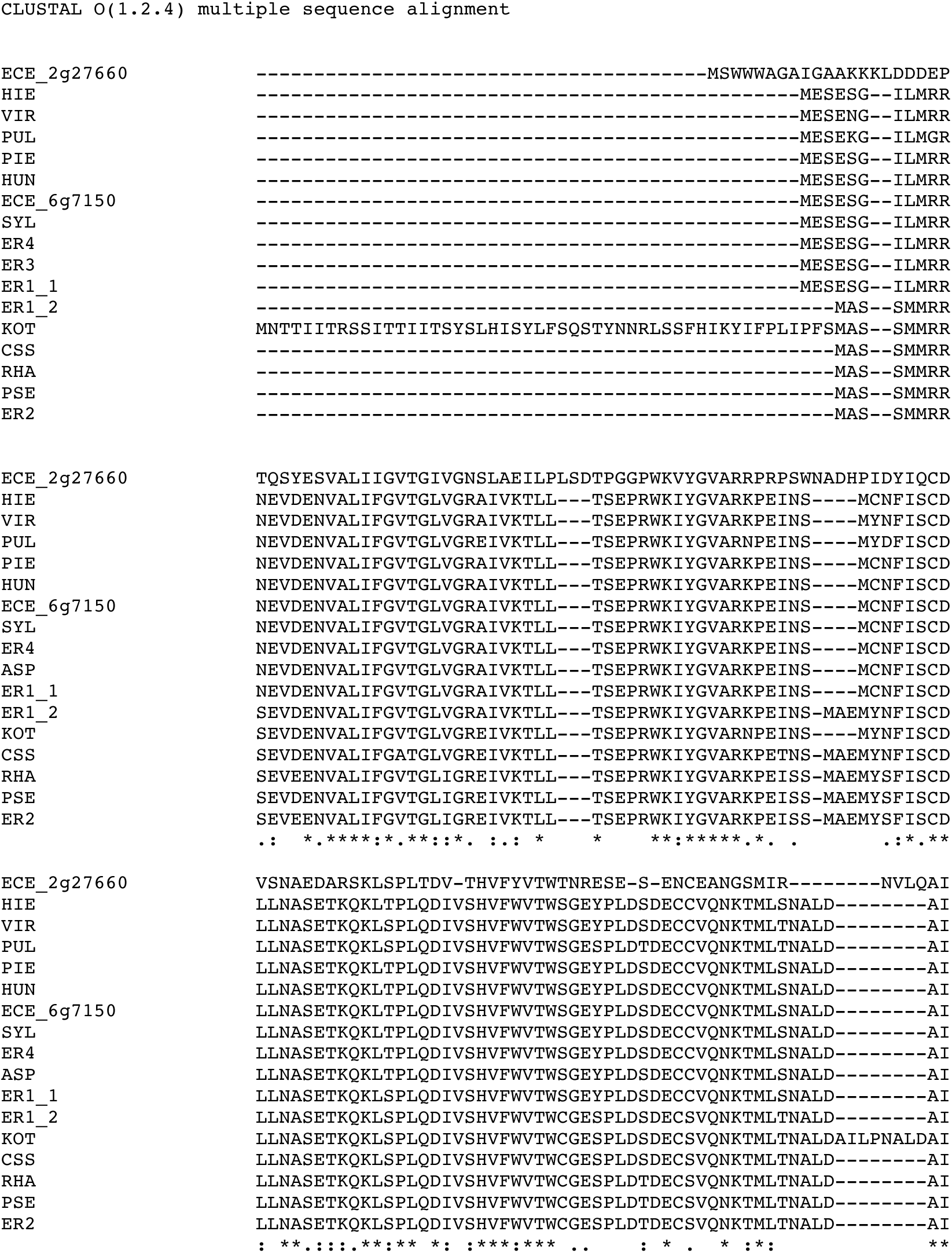
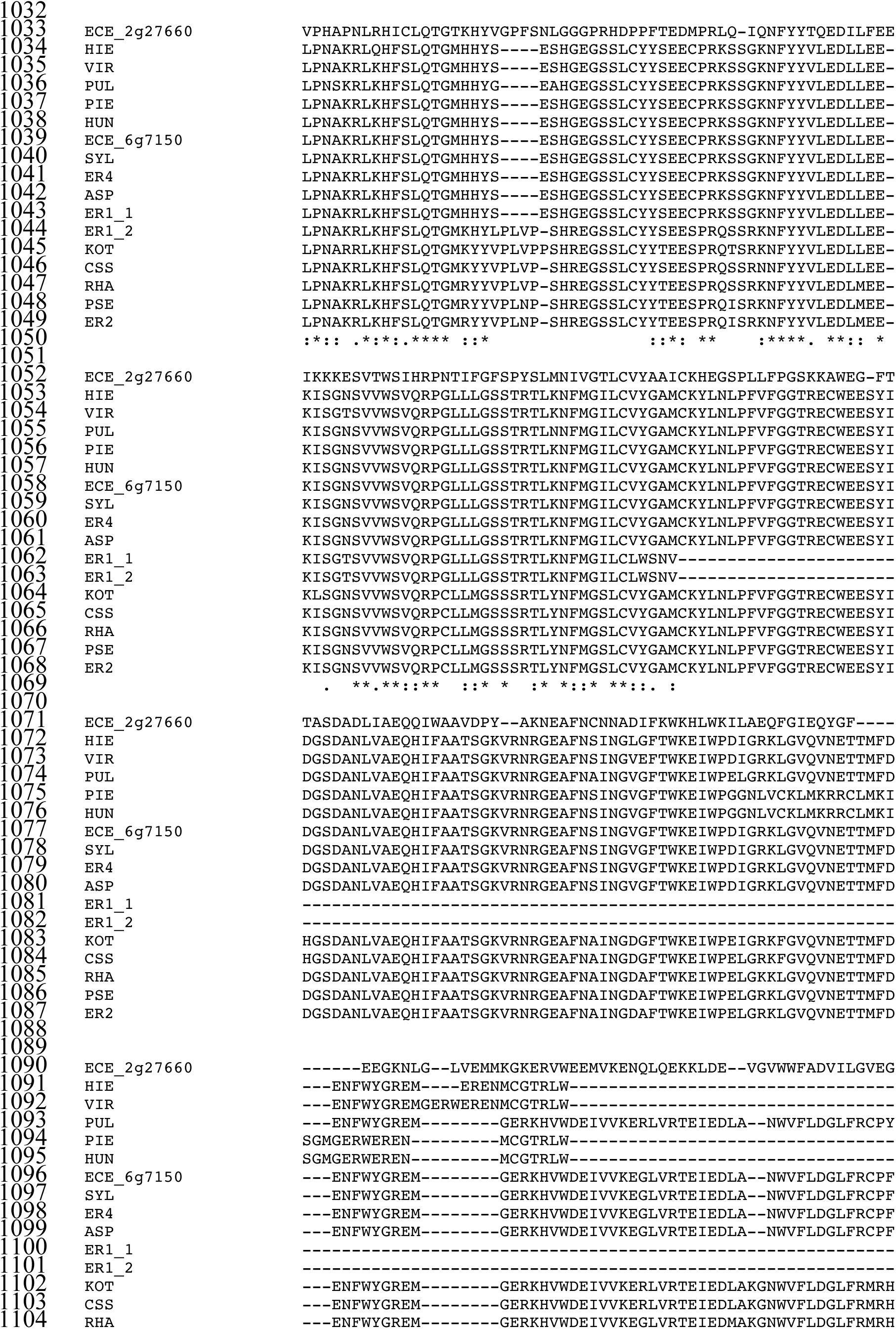
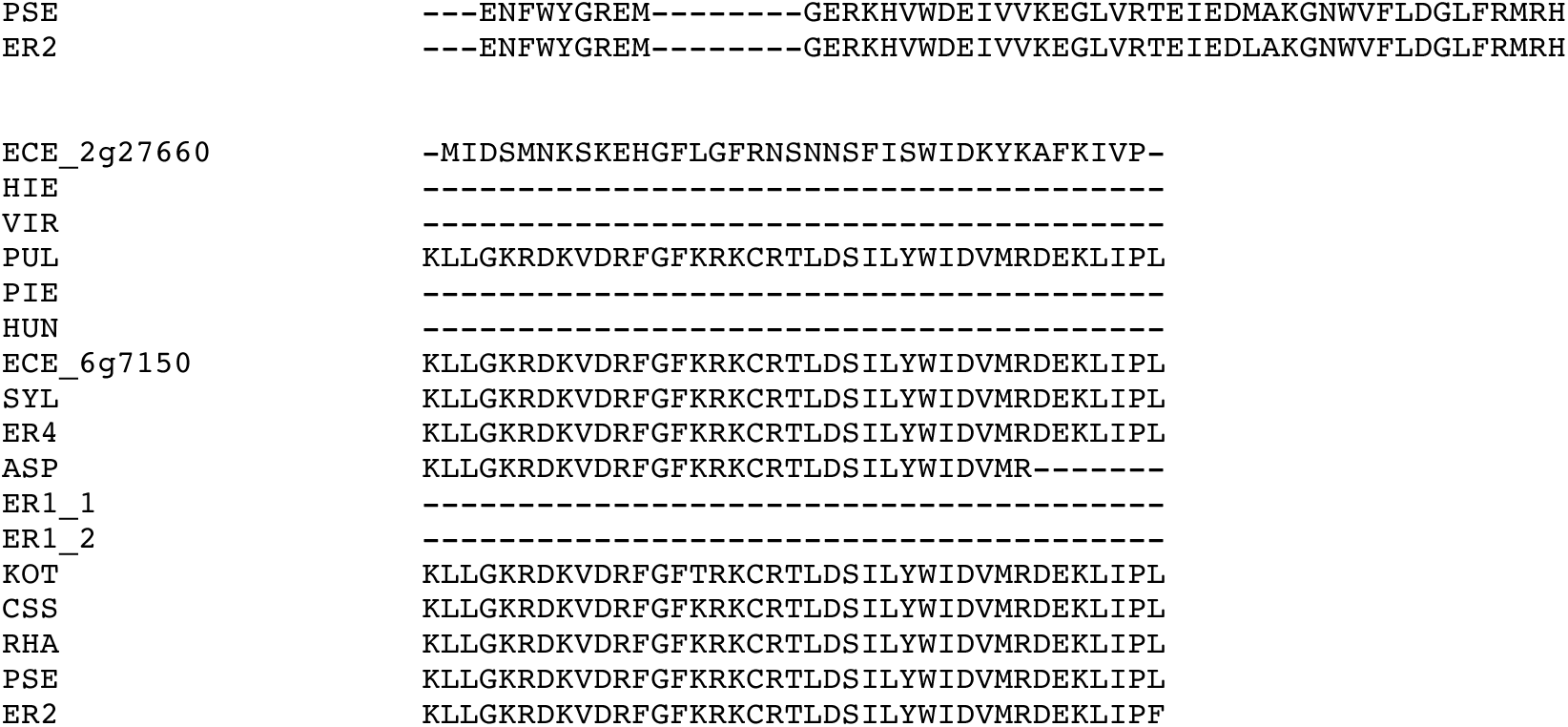
Multiple sequence alignment of progesterone 5β-reductases (P5βR) from *Erysimum* species. All orthologs of *Ec*P5βR2 that could be recovered from transcriptome data are included, and *Ec*P5βR1 (ECE_2g27660) is included as an outgroup. Translated proteins were aligned using Clustal Omega. Species included: *E. crassipes* (CSS), *E. cheiranthoides* (ECE), *Erysimum* sp. 1 (ER1), *Erysimum* sp. 2 (ER2), *Erysimum* sp. 3 (ER3), *Erysimum* sp. 4 (ER4), *E. hieraciifolium* (HIE), *E. hungaricum* (HUN), *E. kotschyanum* (KOT), *E. pieninicum* (PIE), *E. pseudorhaeticum* (PSE), *E. pulchellum* (PUL), *E. rhaeticum* (RHA), *E. sylvestre* (SYL), *E. virgatum* (VIR). Sequences correspond to protein phylogeny in Figure 5c.

**Figure S15.**
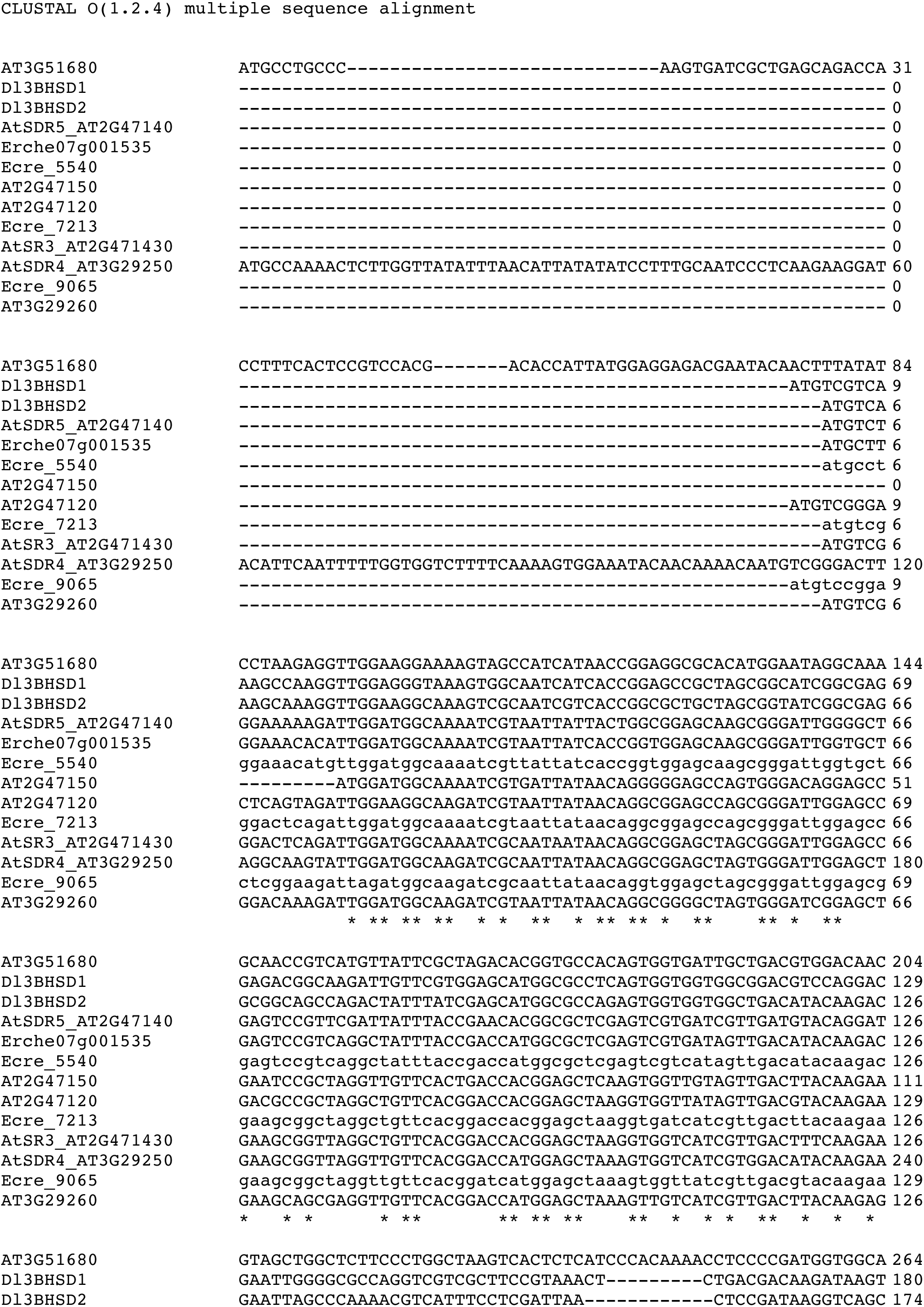
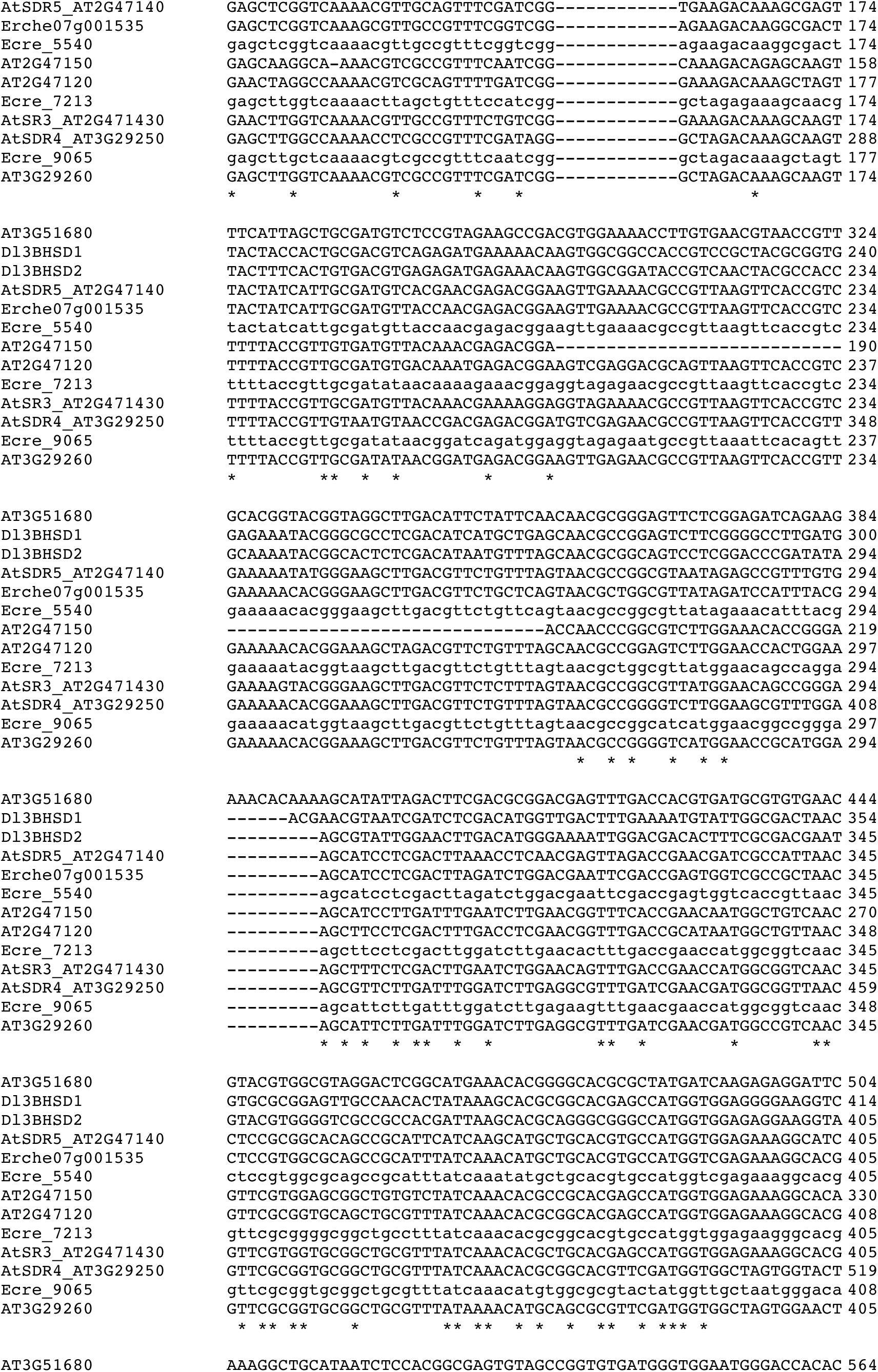
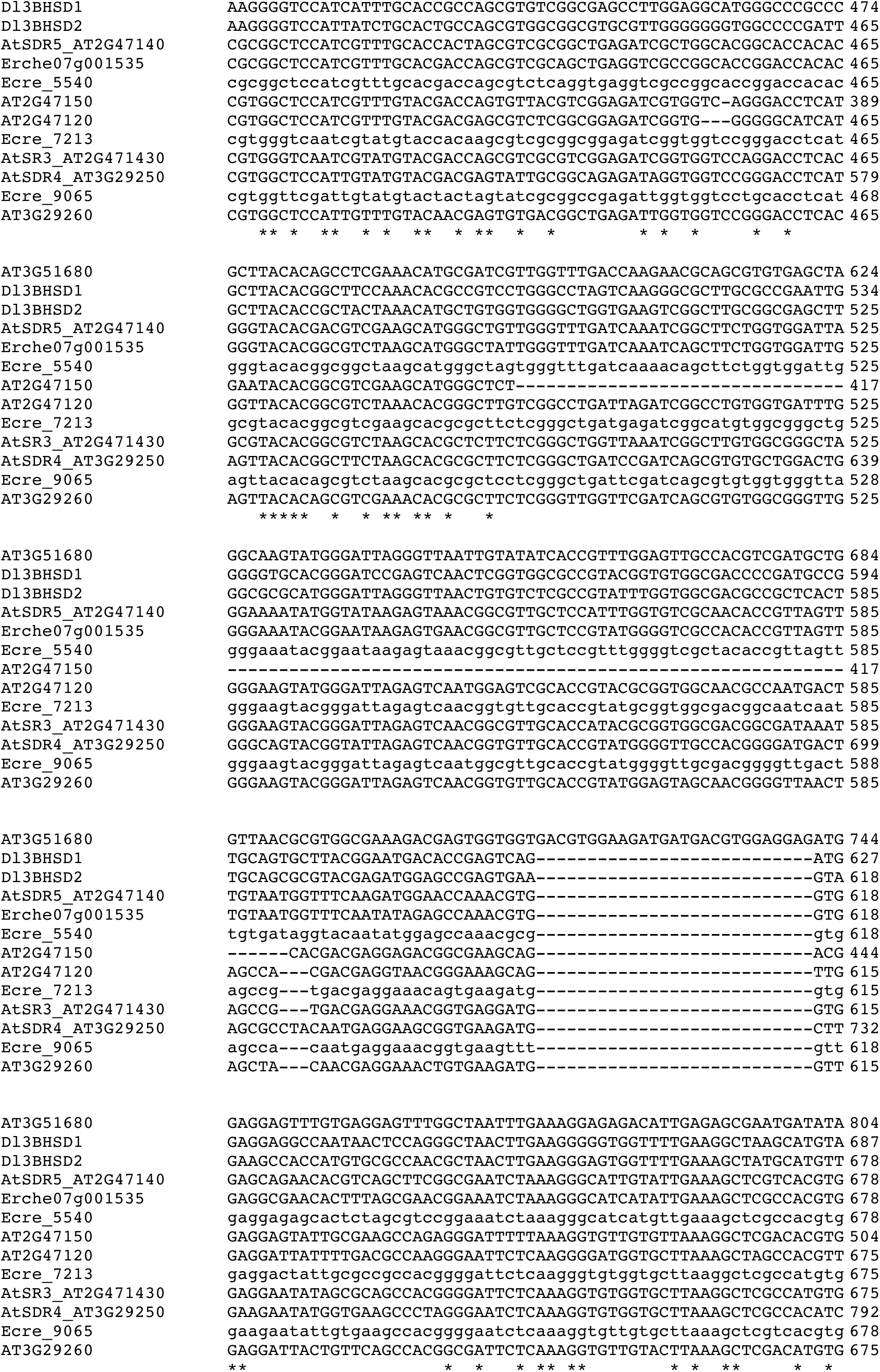
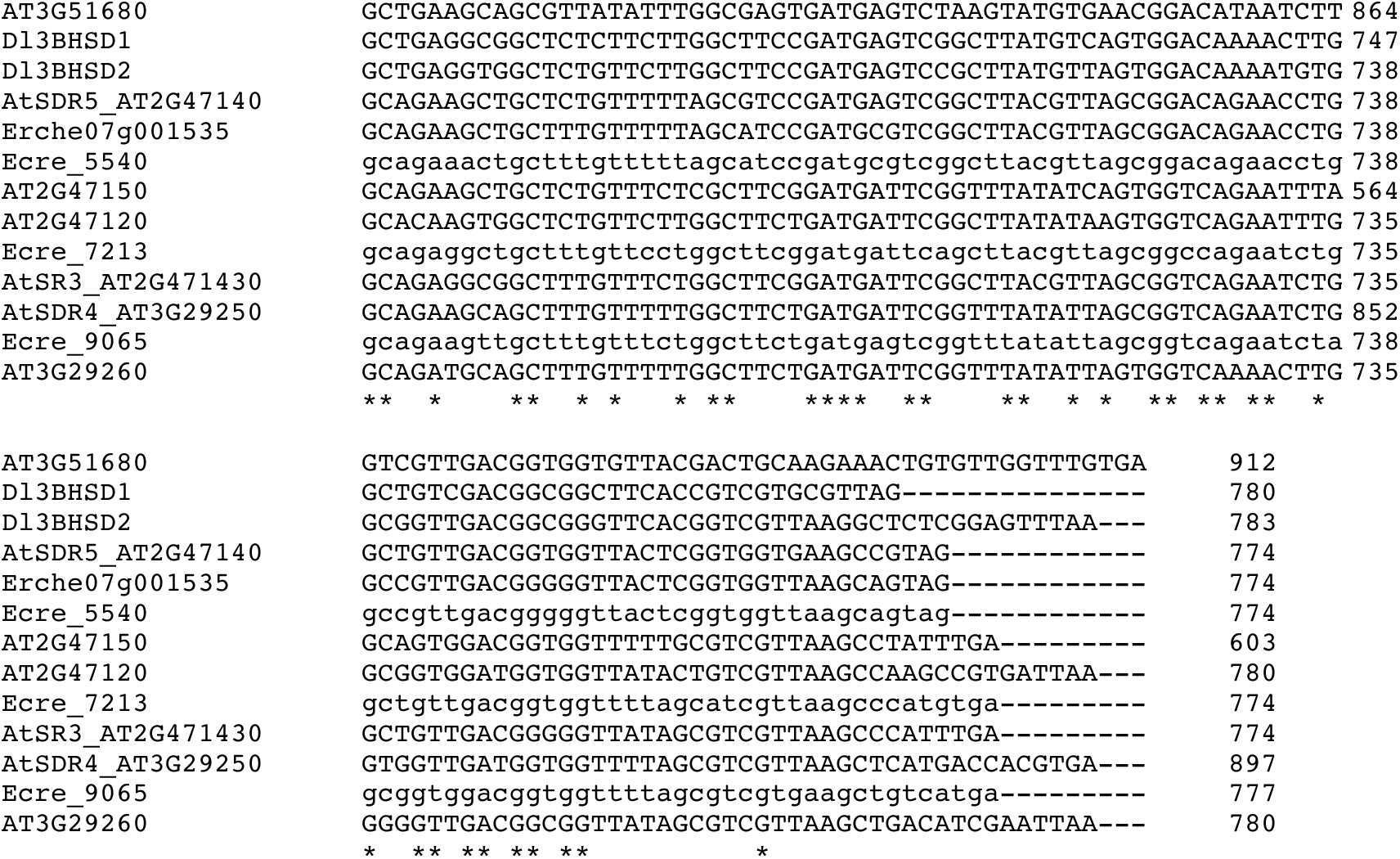
Multiple sequence alignment of 3β-hydroxysteroid dehydrogenases (3βHSD). Coding sequences were aligned using Clustal Omega. Species included: *Arabidopsis thaliana* (AT/At), *Erysimum cheiranthoides* (*Ec*/Erche), *Erysimum crepidifolium* (*Ecre*), and *Digitalis lanata* (*Dl*). Sequences correspond to gene phylogeny in Figure 6a.

**Figure S16.**
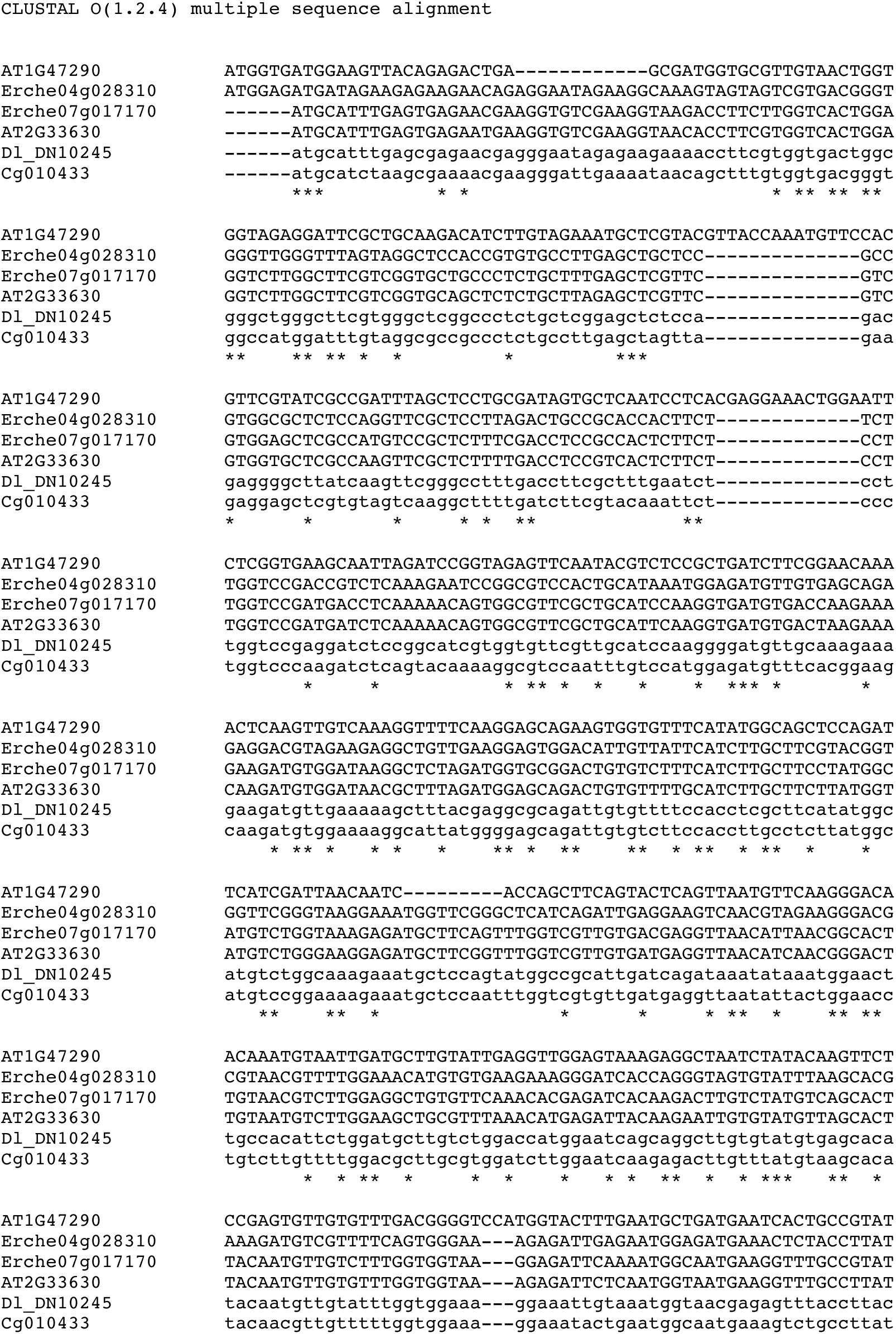
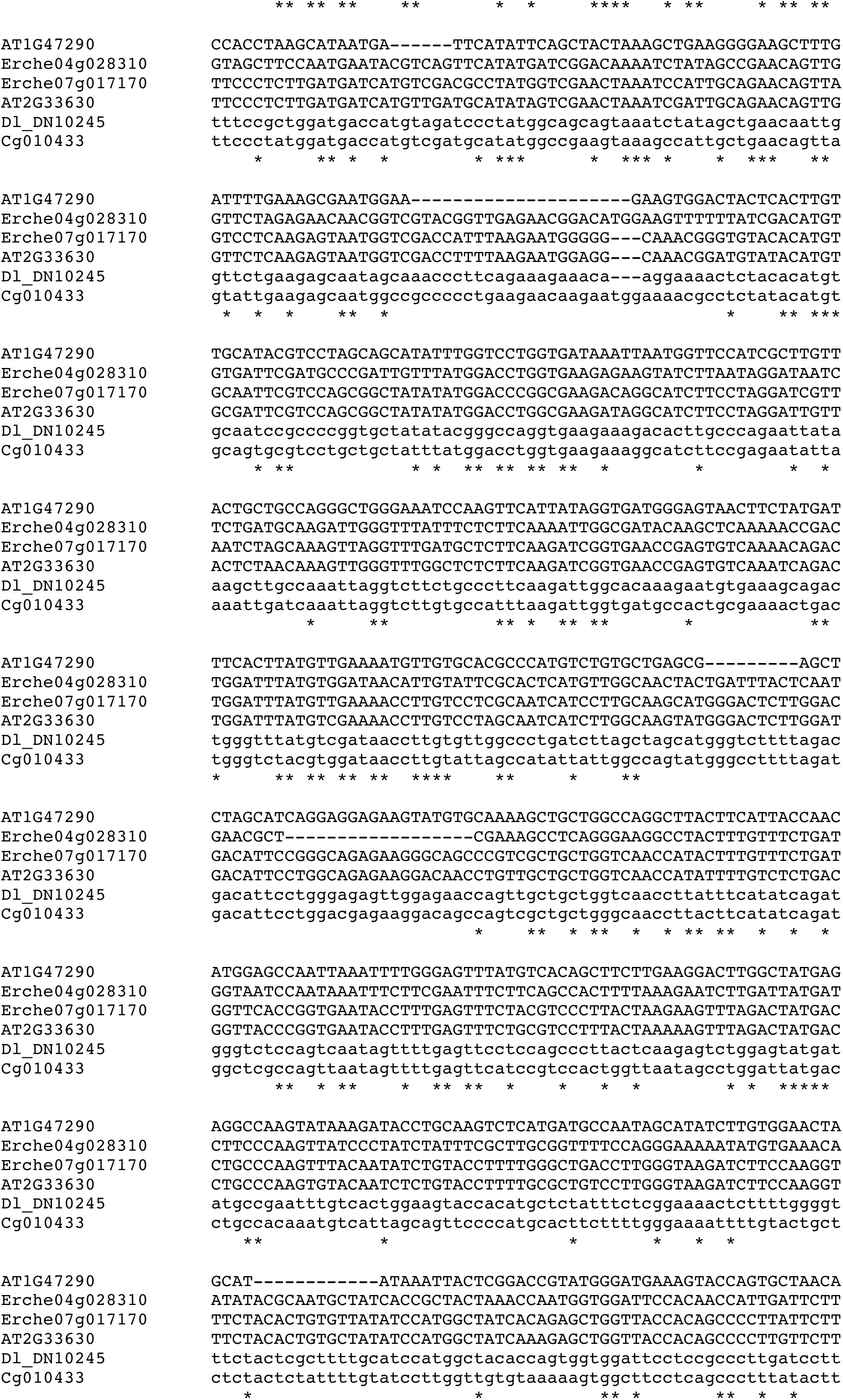
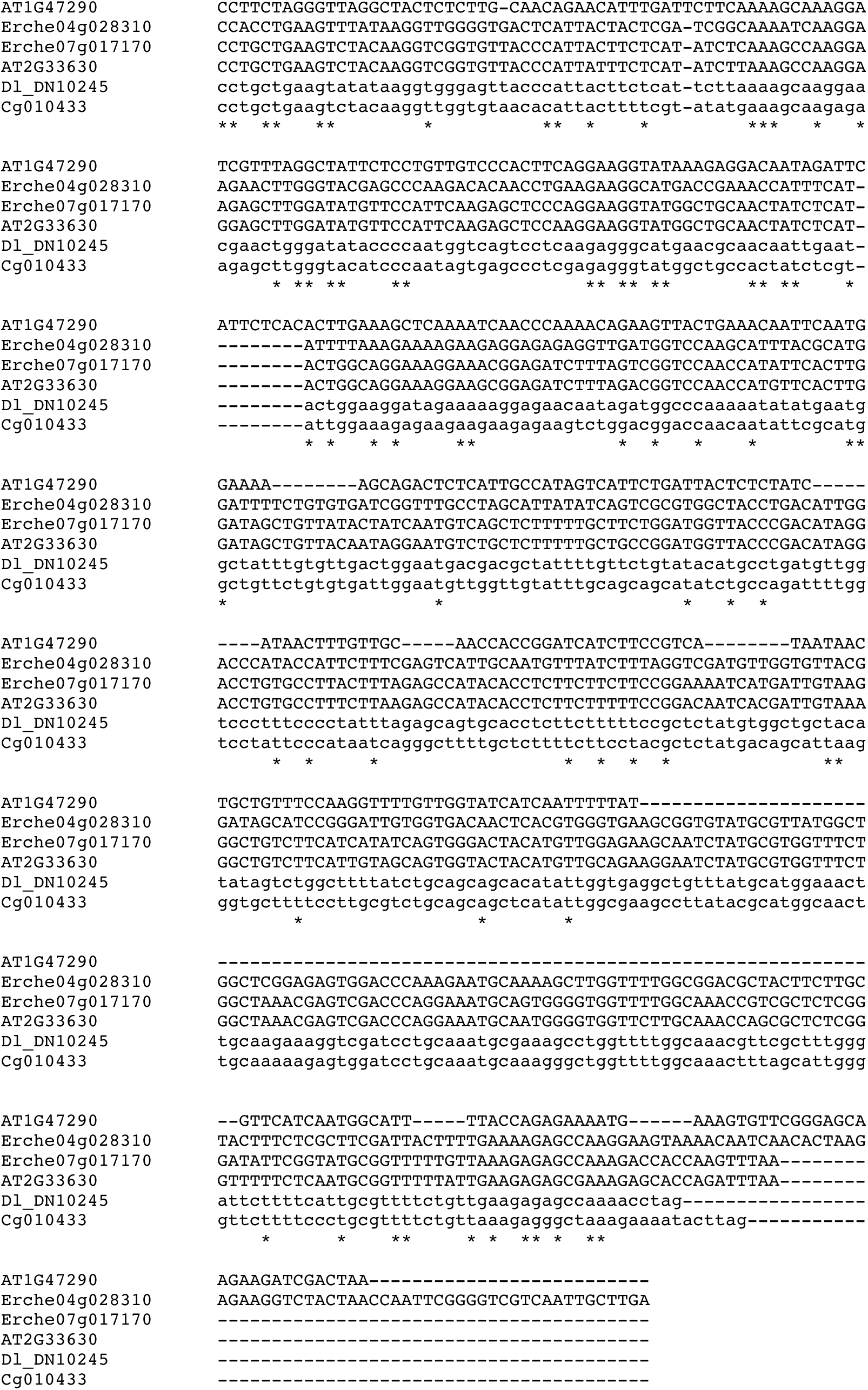
Multiple sequence alignment of 3-ketosteroid isomerases (3KSI). Coding sequences were aligned using Clustal Omega. Species included: *Arabidopsis thaliana* (*At*/AT), *Calotropis gigantea* (*Cg*), *Erysimum cheiranthoides* (*Ec*/Erche), and *Digitalis lanata* (*Dl*). Sequences correspond to gene phylogeny in Figure 6b.

**Figure S17.**
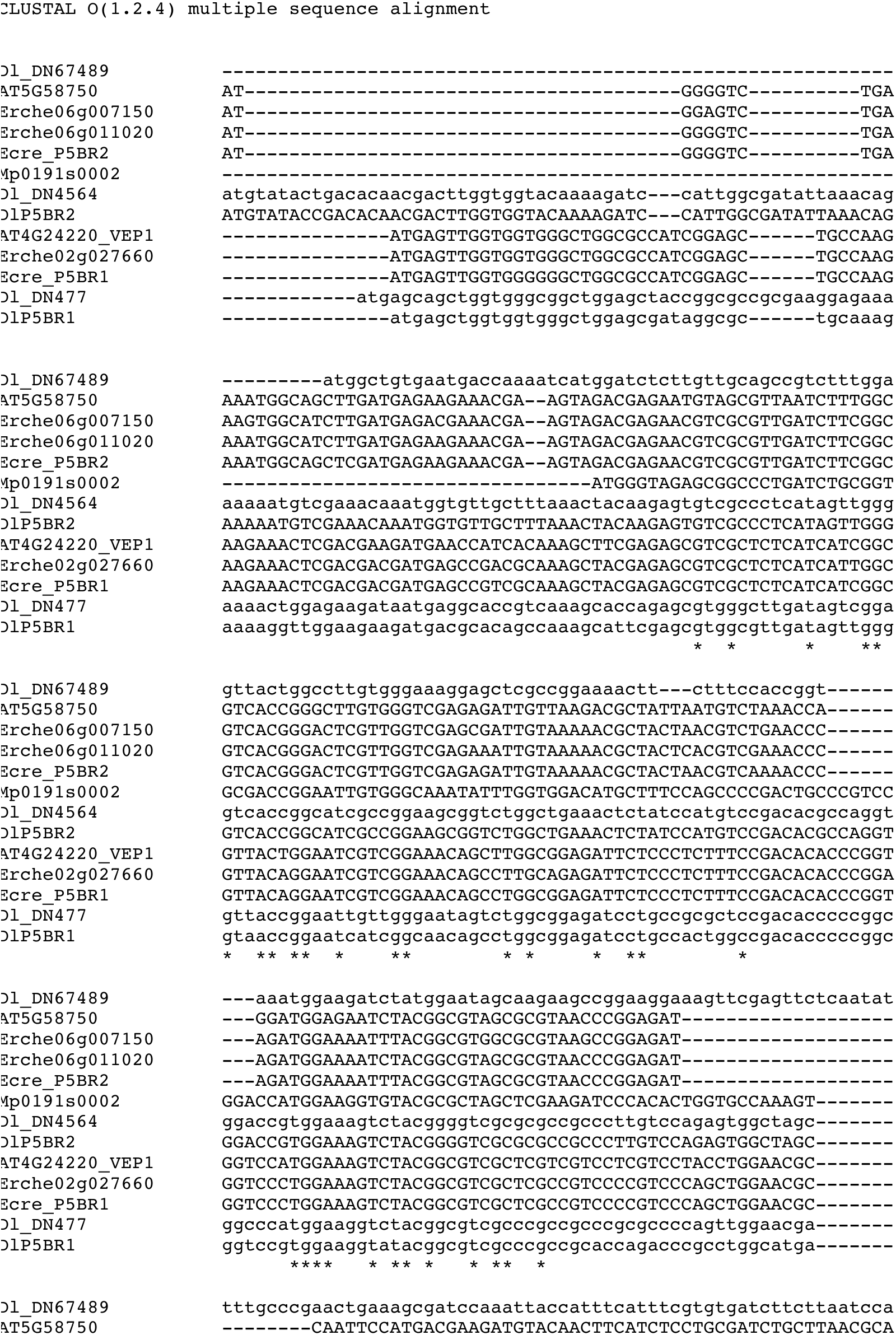
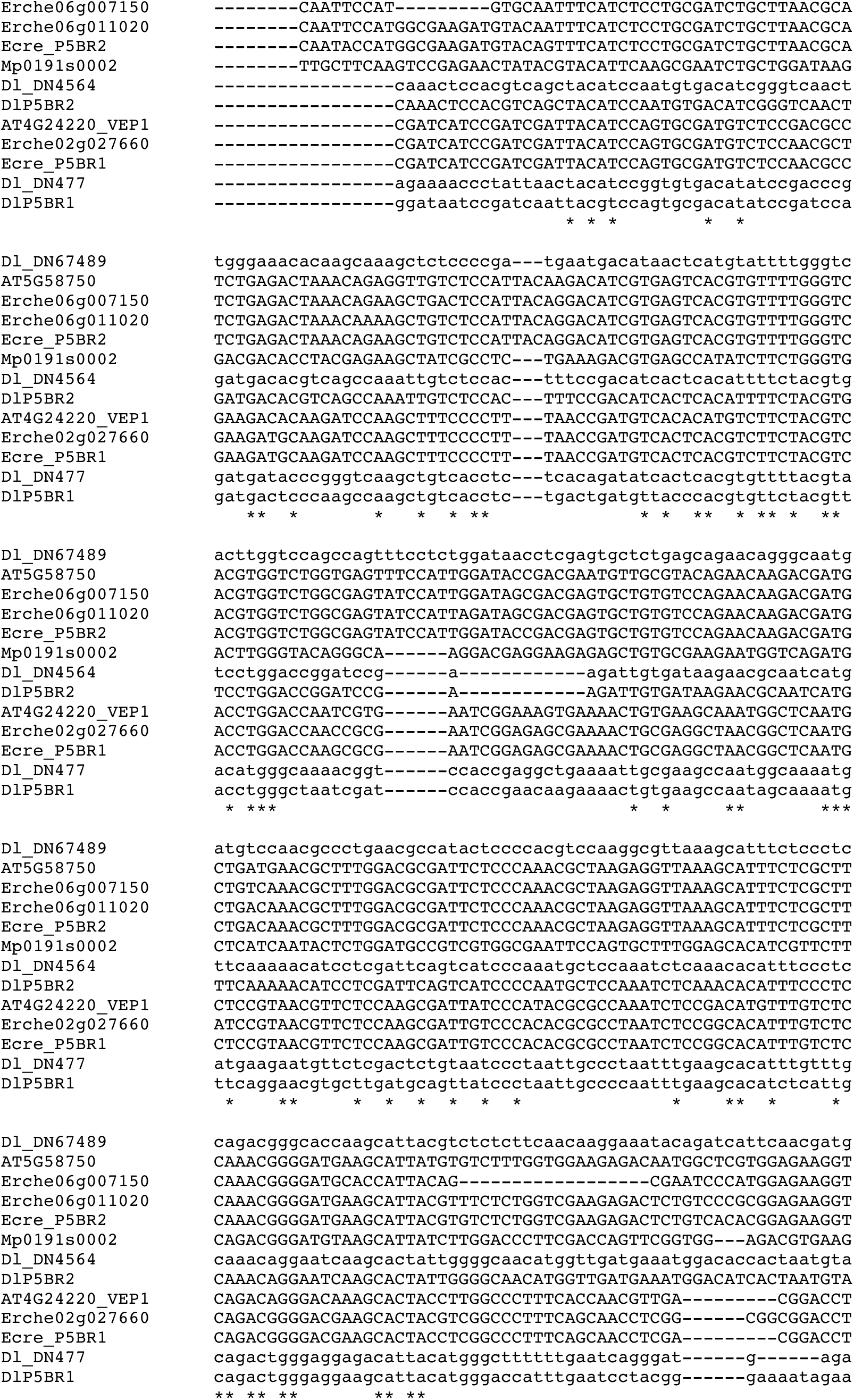
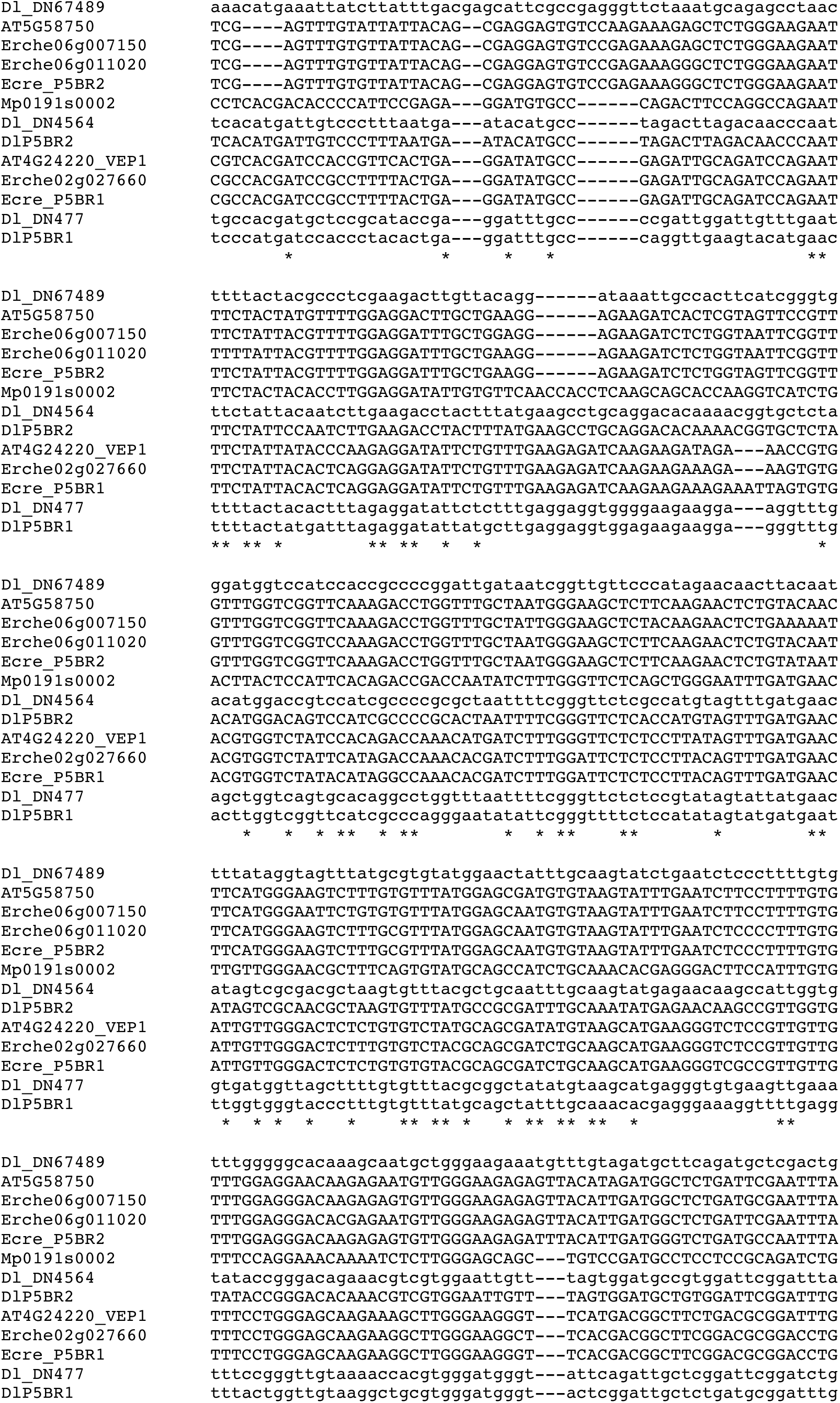
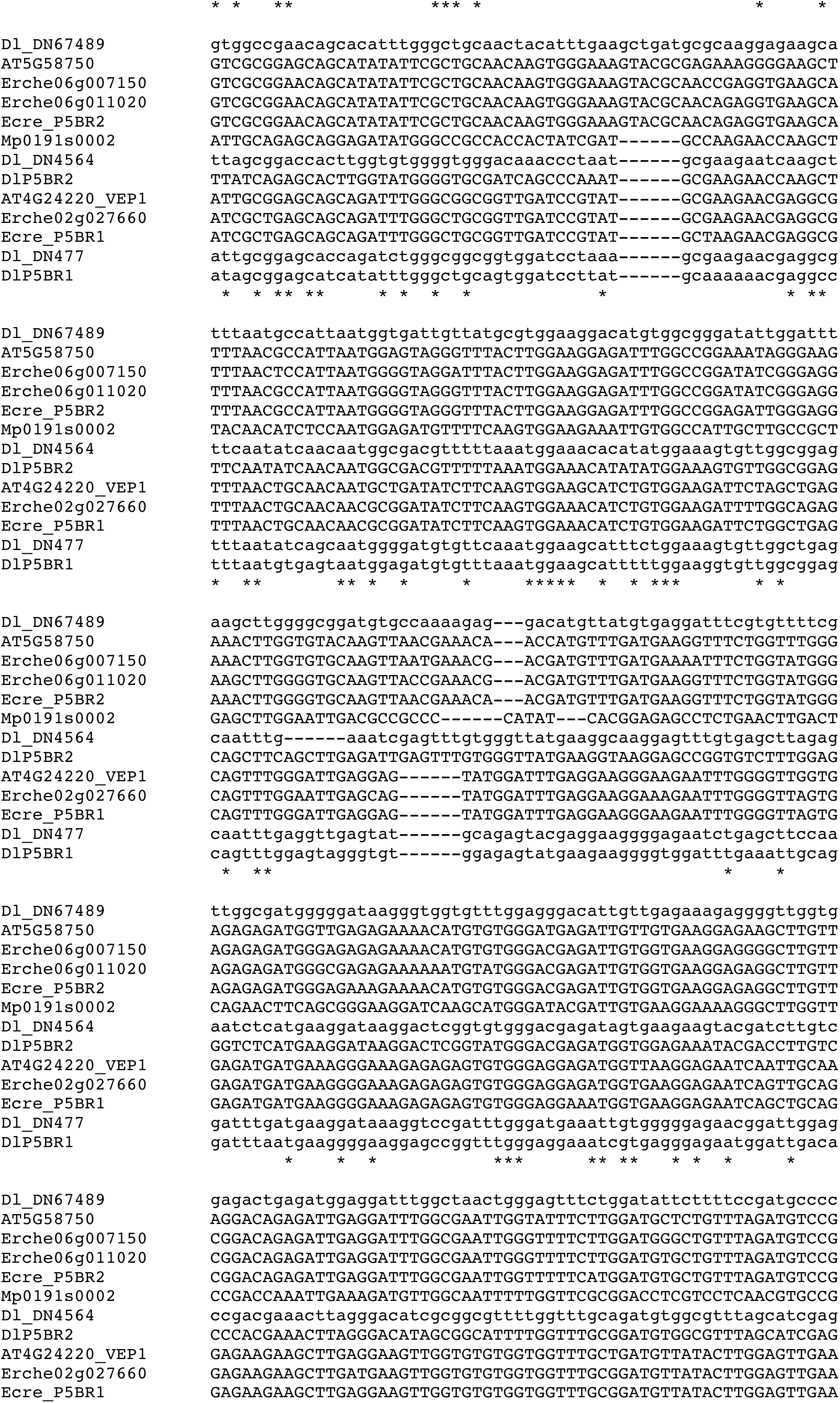
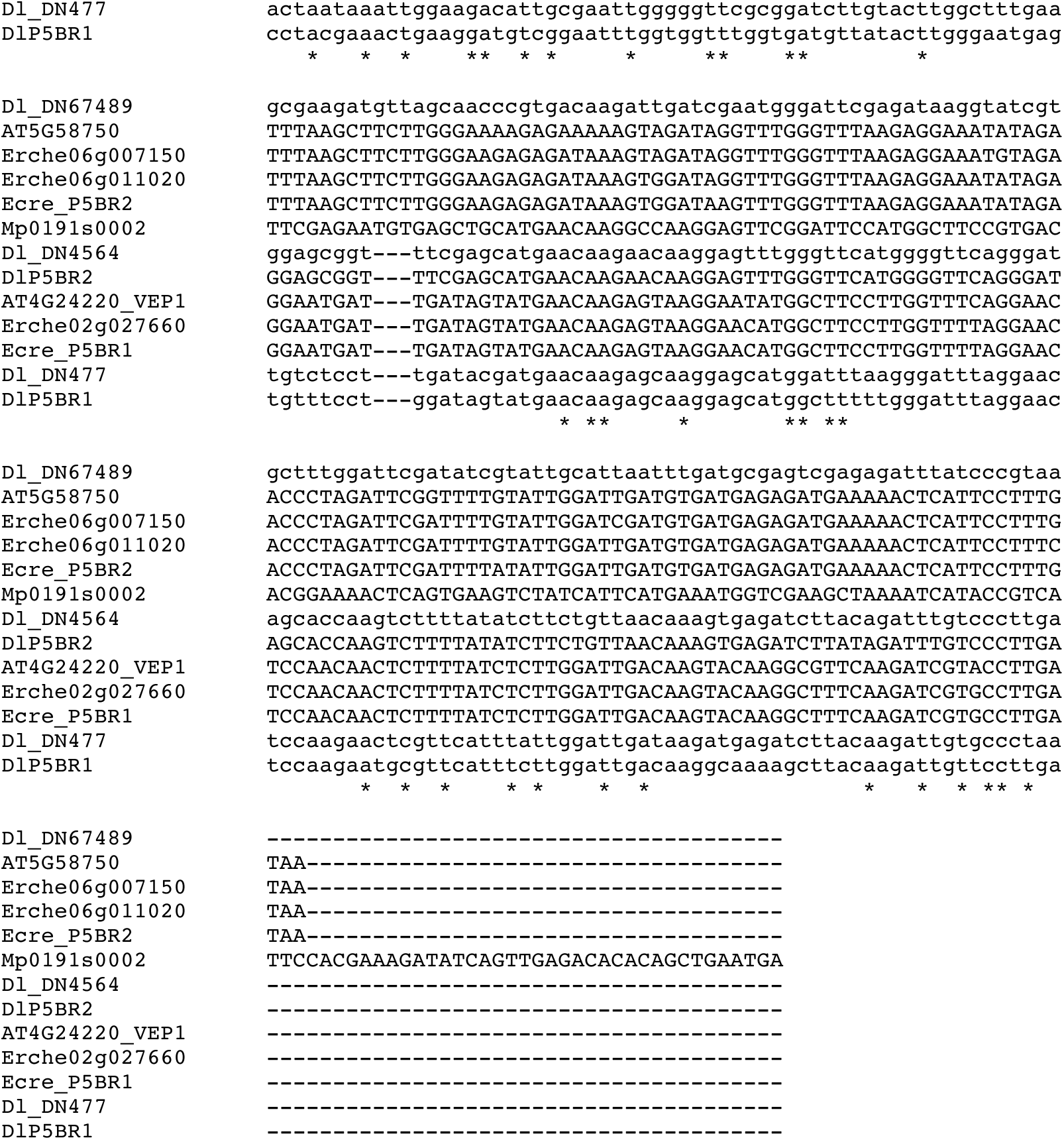
Multiple sequence alignment of Progesterone 5β-reductases (P5βR). Coding sequences were aligned using Clustal Omega. Species included: *Arabidopsis thaliana* (*At*/AT), *Erysimum cheiranthoides* (*Ec*/Erche), *Erysimum crepidifolium* (*Ecre*), *Digitalis lanata* (*Dl*), and *Marchantia polymorpha* (*Mp*). Sequences correspond to gene phylogeny in Figure 6c.

**Figure S18.**
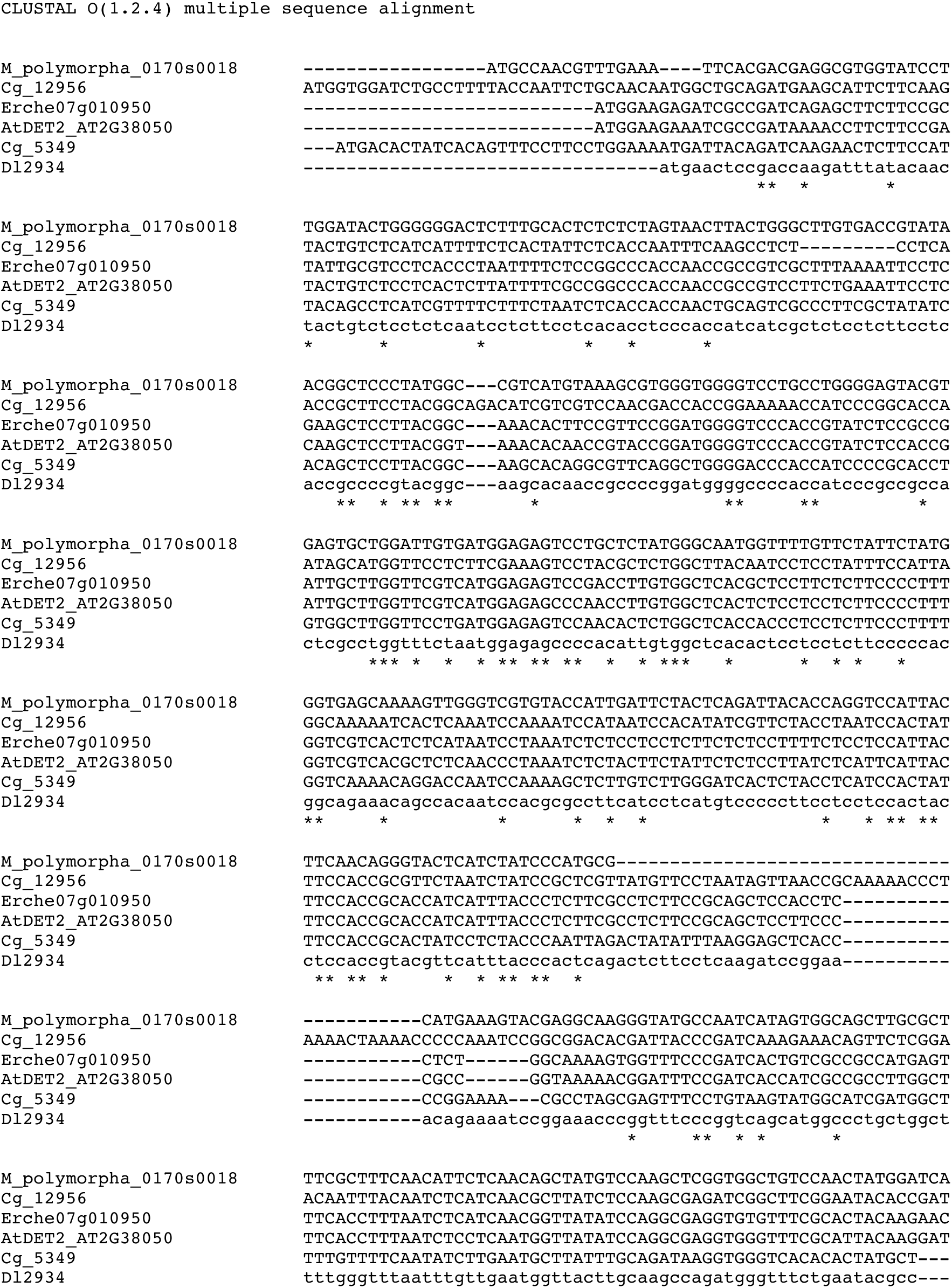
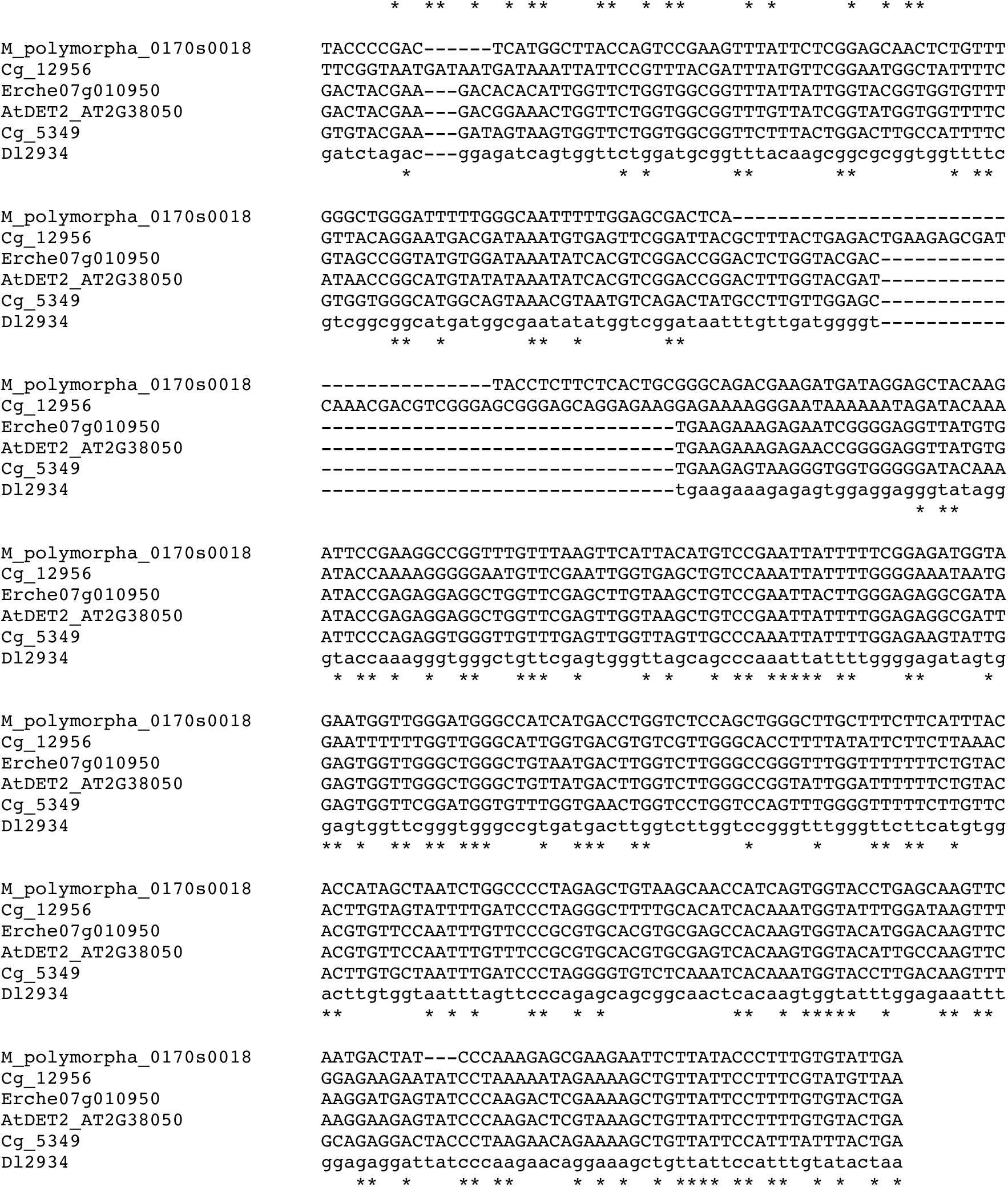
Multiple sequence alignment of steroid 5α-reductases (5αR/DET2). Coding sequences were aligned using Clustal Omega. Species included: *Arabidopsis thaliana* (*At*/AT), *Calotropis gigantea* (*Cg*), *Erysimum cheiranthoides* (*Ec*/Erche), *Digitalis lanata* (*Dl*), and *Marchantia polymorpha* (*Mp*). Sequences correspond to gene phylogeny in Figure 6d.

## Notes

### Competing Interest Statement

The authors have declared no competing interest.

## References

1. G. Fraenkel, The Raison d’Etre of Secondary Plant Substances. Science (80-.). 129, 1466–1470 (1959).

2. P. R. Ehrlich, P. H. Raven, Butterflies and Plants : A Study in Coevolution. Evolution (N. Y). 18, 586–608 (1964).

3. P. Feeny, Defensive Ecology of the Cruciferae. Ann. Missouri Bot. Gard. 64, 221–234 (1977).

4. H. V. Cornell, B. A. Hawkins, Herbivore responses to plant secondary compounds: A test of phytochemical coevolution theory. Am. Nat. 161, 507–522 (2003).

5. M. Berenbaum, Toxicity of a furanocoumarin to armyworms: A case of biosynthetic escape from insect herbivores. Science (80-.). 201, 532–534 (1978).

6. D. L. Forrister, et al., Diversity and divergence: evolution of secondary metabolism in the tropical tree genus Inga. New Phytol. 237, 631–642 (2023).

7. A. A. Agrawal, G. Petschenka, R. A. Bingham, M. G. Weber, S. Rasmann, Toxic cardenolides: Chemical ecology and coevolution of specialized plant-herbivore interactions. New Phytol. 194, 28–45 (2012).

8. W. L. Miller, R. J. Auchus, The molecular biology, biochemistry, and physiology of human steroidogenesis and its disorders. Endocr. Rev. 32, 81–151 (2011).

9. T. J. Bach, M. Rohmer, Isoprenoid Synthesis in Plants and Microorganisms (2013).

10. P. Lindemann, Steroidogenesis in plants - Biosynthesis and conversions of progesterone and other pregnane derivatives. Steroids 103, 145–152 (2015).

11. J. Munkert, et al., Progesterone 5β-reductase genes of the Brassicaceae family as function-associated molecular markers. Plant Biol. 17, 1113–1122 (2015).

12. K. Schmidt, et al., PRISEs (progesteron 5b-reductase and/or iridoid synthase-like 1,4-enone reductases): Catalytic and substrate promiscuity allows for realization of multiple pathways in plant metabolism. Phytochemistry 156, 9–19 (2018).

13. V. Herl, G. Fischer, F. Mu, W. Kreis, Molecular cloning and heterologous expression of progesterone 5b-reductase from *Digitalis lanata* Ehrh. Phytochemistry 67, 225–231 (2006).

14. W. Kreis, The Foxgloves (Digitalis) Revisited. Planta Med. 83, 962–976 (2017).

15. S. Wen, et al., Cardenolides from the Apocynaceae family and their anticancer activity. Fitoterapia 112, 74–84 (2016).

16. H. El-Askary, J. Hölzl, S. Hilal, E. S. El-Kashoury, A comparative study of the cardenolide content of different organs of Gomphocarpus sinaicus. Phytochemistry 38, 1181–1184 (1995).

17. T. Warashina, K. Shikata, T. Miyase, S. Fujii, T. Noro, New cardenolide and acylated lignan glycosides from the aerial parts of Asclepias curassavica. Chem. Pharm. Bull. 56, 1159–1163 (2008).

18. J. Xu, K. Takeya, H. Itokawa, Pregnanes and cardenolides from Periploca sepium. Phytochemistry 29, 344–346 (1990).

19. F. Schaller, W. Kreis, Cardenolide genin pattern in Isoplexis plants and shoot cultures. Planta Med. 72, 1149–1156 (2006).

20. S. Spengel, E. Hauser, H. H. Linde, A. X. Vaz, K. Meyer, Die Glykoside der Blstter von Isoplexis canariensis (L.) G. Don. Helv. Chim. Acta 50, 1893–1911 (1967).

21. V. K. Saxena, P. K. Chaturvedi, A novel cardenolide, canarigenin-3-O-α-L-rhamnopyranosyl-(1->5)-O-β-D-xylofuranoside, from the rhizomes of Convallaria majalis. J. Nat. Prod. 55, 39–42 (1992).

22. I. F. Makarevich, K. V. Zhernoklev, T. V. Slyusarskaya, G. N. Yarmolenko, Cardenolide-containing plants of the family Cruciferae. Chem. Nat. Compd. 30, 275–289 (1994).

23. J. A. Moore, C. Tamm, T. Reichstein, Die Glykoside der Goldlacksamen, Cheiranthus Cheiri L. Helv. Chim. Acta 37, 755–770 (1954).

24. I. F. Makarevich, Cheiranthus allioni--A unique cardenolide-bearing plant. Chem. Nat. Compd. 28, 265–271 (1992).

25. J. Nielsen, Host plant selection of monophagous and oligophagous flea beetles feeding on crucifers. Entomol. Exp. Appl. 24, 362–369 (1978).

26. G. Petschenka, et al., Relative selectivity of plant cardenolides for Na+/K+-ATPases from the monarch butterfly and non-resistant insects. Front. Plant Sci. 9, 1–13 (2018).

27. C. Tamm, “New aspects of cardiac glycosides: The stereochemistry of the glycosides in relation to biological activity” in Proceedings of the First International Pharmacological Meeting, B. Uvnās, W. Wilbrandt, P. Lindgren, Eds. (The Macmillan Company, 1963), pp. 11–26.

28. S. Chhajed, et al., Glucosinolate Biosynthesis and the Glucosinolate–Myrosinase System in Plant Defense Shweta. Agronomy 10 (2020).

29. S. Zhou, A. Richter, G. Jander, B. Thompson, T. Road, Beyond Defense : Multiple Functions of Benzoxazinoids in Maize Metabolism Special Focus Issue – Mini Review. Plant Cell Physiol. 59, 1528–1537 (2018).

30. T. Leykauf, et al., Overexpression and RNAi-mediated Knockdown of Two 3β-hydroxy-Δ5-steroid dehydrogenase Genes in *Digitalis lanata* Shoot Cultures Reveal Their Role in Cardenolide Biosynthesis. Planta Med. 89, 833–847 (2023).

31. V. Herl, J. Frankenstein, N. Meitinger, F. Müller-Uri, W. Kreis, Δ5-3β-hydroxysteroid dehydrogenase (3βHSD) from *Digitalis lanata.* Heterologous expression and characterisation of the recombinant enzyme. Planta Med. 73, 704–710 (2007).

32. S. Seidel, W. Kreis, E. Reinhard, Δ5-3β-hydroysteroid dehydrogenase/Δ5-Δ4-ketosteroid isomerase (3β-HSD), a possible enzyme of cardiac glycoside biosynthesis, in cell cultures and plants of Digitalis lanata EHRH. Plant Cell Rep. 8, 621–624 (1990).

33. M. Mirzaei, et al., Less Is More: a Mutation in the Chemical Defense Pathway of Erysimum cheiranthoides (Brassicaceae) Reduces Total Cardenolide Abundance but Increases Resistance to Insect Herbivores. J. Chem. Ecol. 46, 1131–1143 (2020).

34. J. Munkert, et al., Iridoid synthase activity is common among the plant progesterone 5β-reductase family. Mol. Plant 8, 136–152 (2015).

35. G. C. Younkin, et al., Cardiac glycosides protect wormseed wallflower (*Erysimum cheiranthoides*) against some, but not all, glucosinolate-adapted herbivores. New Phytol. (2024) 10.1111/nph.19534.

36. T. Paysan-Lafosse, et al., InterPro in 2022. Nucleic Acids Res. 51, D418–D427 (2023).

37. S. Fujioka, et al., The Arabidopsis deetiolated2 mutant is blocked early in brassinosteroid biosynthesis. Plant Cell 9, 1951–1962 (1997).

38. N. Meitinger, et al., The catalytic mechanism of the 3-ketosteroid isomerase of *Digitalis lanata* involves an intramolecular proton transfer and the activity is not associated with the 3β-hydroxysteroid dehydrogenase activity. Tetrahedron Lett. 57, 1567–1571 (2016).

39. N. Meitinger, D. Geiger, T. W. Augusto, R. Maia De Pádua, W. Kreis, Purification of Δ5-3-ketosteroid isomerase from *Digitalis lanata*. Phytochemistry 109, 6–13 (2015).

40. F. Rosati, et al., 5α-Reductase activity in Lycopersicon esculentum: Cloning and functional characterization of LeDET2 and evidence of the presence of two isoenzymes. J. Steroid Biochem. Mol. Biol. 96, 287–299 (2005).

41. M. Kunert, et al., Promiscuous CYP87A enzyme activity initiates cardenolide biosynthesis in plants. Nat. Plants 9, 1607–1617 (2023).

42. A. Pandey, V. Swarnkar, T. Pandey, P. Srivastava, Transcriptome and Metabolite analysis reveal candidate genes of the cardiac glycoside biosynthetic pathway from Calotropis procera. Sci. Rep., 1–14 (2016).

43. T. Züst, et al., Independent evolution of ancestral and novel defenses in a genus of toxic plants (Erysimum, Brassicaceae). Elife 9, 1–42 (2020).

44. J. Munkert, M. Ernst, F. Müller-Uri, W. Kreis, Identification and stress-induced expression of three 3β-hydroxysteroid dehydrogenases from Erysimum crepidifolium Rchb. and their putative role in cardenolide biosynthesis. Phytochemistry 100, 26–33 (2014).

45. J. Munkert, P. Bauer, E. Burda, F. Müller-Uri, W. Kreis, Progesterone 5β-reductase of *Erysimum crepidifolium*: CDNA cloning, expression in *Escherichia coli*, and reduction of enones with the recombinant protein. Phytochemistry 72, 1710–1717 (2011).

46. P. Bauer, et al., Highly conserved progesterone 5 b-reductase genes (P5bR) from 5b-cardenolide-free and 5b-cardenolide-producing angiosperms. Phytochemistry 71, 1495– 1505 (2010).

47. J. Klein, et al., RNAi-mediated gene knockdown of progesterone 5β-reductases in *Digitalis lanata* reduces 5β-cardenolide content. Plant Cell Rep. 40, 1631–1646 (2021).

48. E. Carroll, B. R. Gopal, I. Raghavan, M. Mukherjee, Z. Q. Wang, A cytochrome P450 CYP87A4 imparts sterol side-chain cleavage in digoxin biosynthesis. Nat. Commun. 14, 4042 (2023).

49. N.-C. Ha, G. Choi, K. Y. Choi, B.-H. Oh, Structure and enzymology of Δ5-3-ketosteroid isomerase. Curr. Opin. Struct. Biol. 11, 674–678 (2001).

50. D. M. Piatak, P.-F. L. Tang, P. D. Sørensen, Constituents of *Erysimum inconspicuum*. Two sulfur-containing lactone compounds. J. Nat. Prod. 48, 424–428 (1985).

51. T. Züst, M. Mirzaei, G. Jander, *Erysimum cheiranthoides*, an ecological research system with potential as a genetic and genomic model for studying cardiac glycoside biosynthesis. Phytochem. Rev. 17, 1239–1251 (2018).

52. M. E. Deluca, A. M. Seldes, E. G. Gros, The 14ß-HydroxyIation in the Biosynthesis of Cardenolides in *Digitalis purpurea*. The Role of 3ß-Hydroxy-5ß-pregn-8(14)-en-20-one. Zeitschrift für Naturforsch. 42c, 77–78 (1987).

53. P. D. Sonawane, et al., Short-chain dehydrogenase/reductase governs steroidal specialized metabolites structural diversity and toxicity in the genus Solanum. Proc. Natl. Acad. Sci. 115, E5419–E5428 (2018).

54. K. Repke, New developments in cardiac glycoside structure-activity relationships. Trends Pharmacol. Sci. 6, 275–278 (1982).

55. S. R. Strickler, A. F. Powell, L. A. Mueller, T. Zust, G. Jander, NCBI BioProject ID PRJNA563696. Rapid and independent evolution of ancestral and novel chemical defenses in a genus of toxic plants (*Erysimum*, Brassicaceae) (2019).

56. M. Mirzaei, et al., Aphid resistance segregates independently of cardiac glycoside and glucosinolate content in an *Erysimum cheiranthoides* (wormseed wallflower) F2 population. bioRxiv (2024) 10.1101/2024.01.11.575310.

57. N. L. Bray, H. Pimentel, P. Melsted, L. Pachter, Near-optimal probabilistic RNA-seq quantification. Nat. Biotechnol. 34, 525–527 (2016).

58. J. H. Wisecaver, et al., A global coexpression network approach for connecting genes to specialized metabolic pathways in plants. Plant Cell 29, 944–959 (2017).

59. Terrific Broth (TB) Medium (2015) 10.1101/pdb.rec085894.

60. R Core Team, R: A Language and Environment for Statistical Computing (2020).

61. L. Gatto, K. Lilley, MSnbase - an R/Bioconductor package for isobaric tagged mass spectrometry data visualization, processing and quantitation. Bioinformatics 28, 288–289 (2012).

62. L. Gatto, S. Gibb, J. Rainer, MSnbase, efficient and elegant R-based processing and visualisation of raw mass spectrometry data. bioRxiv(2020).

63. S. Graves, H.-P. Piepho, L. S. with help from Sundar Dorai-Raj, multcompView: Visualizations of Paired Comparisons (2023).

64. R. Kolde, pheatmap: Pretty Heatmaps (2019).

65. T. Z. Berardini, et al., The Arabidopsis information resource: Making and mining the “gold standard” annotated reference plant genome. Genesis 53, 474–485 (2015).

66. G. M. Hoopes, et al., Genome assembly and annotation of the medicinal plant *Calotropis gigantea* a producer of anticancer and antimalarial cardenolides. G3 Genes, Genomes, Genet. 8, 385–391 (2018).

67. M. K. Tello-Ruiz, P. Jaiswal, D. Ware, “Gramene: A Resource for Comparative Analysis of Plants Genomes and Pathways” in Plant Bioinformatics, D. Edwards, Ed. (Springer US, 2022), pp. 101–131.

68. F. Sievers, et al., Fast, scalable generation of high-quality protein multiple sequence alignments using Clustal Omega. Mol. Syst. Biol. 7 (2011).

69. F. Madeira, et al., Search and sequence analysis tools services from EMBL-EBI in 2022. Nucleic Acids Res. 50, W276–W279 (2022).

70. B. Q. Minh, et al., IQ-TREE 2: New Models and Efficient Methods for Phylogenetic Inference in the Genomic Era. Mol. Biol. Evol. 37, 1530–1534 (2020).

71. J. Trifinopoulos, L. T. Nguyen, A. von Haeseler, B. Q. Minh, W-IQ-TREE: a fast online phylogenetic tool for maximum likelihood analysis. Nucleic Acids Res. 44, W232–W235 (2016).

72. D. T. Hoang, O. Chernomor, A. Von Haeseler, B. Q. Minh, L. S. Vinh, UFBoot2: Improving the ultrafast bootstrap approximation. Mol. Biol. Evol. 35, 518–522 (2018).

